# Drugs of abuse hijack a mesolimbic pathway that processes homeostatic need

**DOI:** 10.1101/2023.09.03.556059

**Authors:** Bowen Tan, Caleb J. Browne, Tobias Nöbauer, Alipasha Vaziri, Jeffrey M. Friedman, Eric J. Nestler

**Affiliations:** Laboratory of Molecular Genetics, Howard Hughes Medical Institute, The Rockefeller University, New York, NY 10065, USA; Nash Family Department of Neuroscience, Friedman Brain Institute, Icahn School of Medicine at Mount Sinai, New York, NY 10029, USA; Laboratory of Neurotechnology and Biophysics, The Rockefeller University, New York, NY 10065, USA; The Kavli Neural Systems Institute, The Rockefeller University, New York, NY 10065, USA

## Abstract

Addiction prioritizes drug use over innate needs by “hijacking” brain circuits that direct motivation, but how this develops remains unclear. Using whole-brain FOS mapping and *in vivo* single-neuron calcium imaging, we find that drugs of abuse augment ensemble activity in the nucleus accumbens (NAc) and disorganize overlapping ensemble responses to natural rewards in a cell-type-specific manner. Combining “FOS-Seq”, CRISPR-perturbations, and snRNA-seq, we identify *Rheb* as a shared molecular substrate that regulates cell-type-specific signal transductions in NAc while enabling drugs to suppress natural reward responses. Retrograde circuit mapping pinpoints orbitofrontal cortex which, upon activation, mirrors drug effects on innate needs. These findings deconstruct the dynamic, molecular, and circuit basis of a common reward circuit, wherein drug value is scaled to promote drug-seeking over other, normative goals.

## Introduction

Substance use takes a devastating toll on individuals, their families, and the healthcare system as a whole. As mortality rates continue to rise, there is an urgent need to better understand the pathogenesis of addiction and develop more effective treatments. Abundant research suggests that drugs of abuse cause addiction by usurping brain reward circuits that have been evolutionarily established to direct behavior toward the satisfaction of need states such as hunger or thirst (*1*). Drug-induced changes in the function of these circuits narrows the scope of motivation toward drugs and away from alternative, healthy goals (*2, 3*). Numerous theories of addiction development and maintenance depend on this idea (*4–6*), and further imply that innate neural functions that normally process natural rewards are corrupted by drugs of abuse. However, neurobiological relationships between drug and natural rewards are typically inferred across studies and in separate experimental subjects, thus leaving the underlying physiological and molecular mechanisms linking these functions unclear. To address this gap, we have compared the response of key reward circuits activated by hunger and thirst to the response to morphine and cocaine in the same animals.

Here, we outline a neural substrate that enables drugs of abuse to access, augment, and corrupt a shared pathway that normally subserves physiological needs, ultimately disrupting goal-directed behavior. Unbiased, brain-wide activity mapping identified the nucleus accumbens (NAc), a well-established node for motivation (*7*), as a common hub for cocaine and morphine action. We then used two-photon calcium imaging to track individual neural responses across natural and drug rewards to outline cell-type-specific ensemble dynamics that integrate these reward classes. Combining *in silico* “FOS-seq”, CRISPR-perturbations, and snRNA-seq, we demonstrate that *Rheb*, an mTOR signaling partner, is a crucial molecular substrate that enables drugs to gain access to neurons that process natural reward. Single-cell RNAseq revealed further that *Rheb* KO may contribute to cell-type-specific divergent effects of cocaine and morphine within the NAc. Finally, combining whole-brain activity mapping with retrograde tracing from the NAc identified key input structures that are commonly activated by cocaine and morphine. These higher-order nodes point to a circuit-dependent reward scaling mechanism that relays values of both natural and drug rewards. These results address questions about the circuit architecture, neural coding principles, and molecular restructuring that enable drugs to overtake brain reward circuits (*8*), and point to cell-type-specific mTOR pathways as potential targets for treatment of addiction.

## Results

### NAc is a central integrator of drug and natural reward

To establish a relationship between natural and drug reward processing, we determined how cocaine (a psychostimulant) and morphine (an opioid) affect behavioral responses to hunger and thirst (*9*). We first examined the effects acute cocaine or morphine on feeding and drinking in fasted or dehydrated mice, using doses known to be rewarding (*10*) (Fig. S1a). Fasted mice that received either cocaine or morphine consumed less food during the first 30 min of refeeding compared to saline-treated mice, while morphine exerted a prolonged suppressive effect on food intake that lasted 4 hours post-refeeding (Fig. S1b). In water deprived mice, acute cocaine or morphine exposure likewise decreased water intake during the 30-min rehydration period compared to saline, which recovered within 2-4 hours (Fig. S1b).

We next measured the effects of repeated drug exposure on ad libitum food or water intake over the course of a five-day, daily treatment regimen. Repeated exposure to either drug significantly reduced food intake, water intake, as well as body weight compared to saline treatment (Fig. 1a, b). We tested whether withdrawal from repeated exposure to cocaine or morphine also affects behavioral responses to hunger and thirst. Mice were treated with cocaine, morphine or saline for 5 days, followed by saline administration for additional 3 days to produce spontaneous withdrawal (Fig. S1c). During this 3-day withdrawal, mice were fasted or water deprived overnight and then introduced into a new cage where they were provided free access to food for 20 min or water for 5 min. Mice with prior chronic exposure to cocaine or morphine consumed less food or water compared to mice chronically treated with saline (Fig. S1d). We also performed a sucrose preference test to assess anhedonic-like responses in withdrawal and found that, while mice treated with cocaine showed normal sucrose preference, morphine-treated mice exhibited reductions in sucrose preference (Fig. S1d). These results show that both acute and repeated administration of two drugs of abuse interferes with natural reward consumption, in both ad libitum conditions or after deprivation, and that these effects persist during withdrawal when drug is no longer on board.

**Figure. 1.**
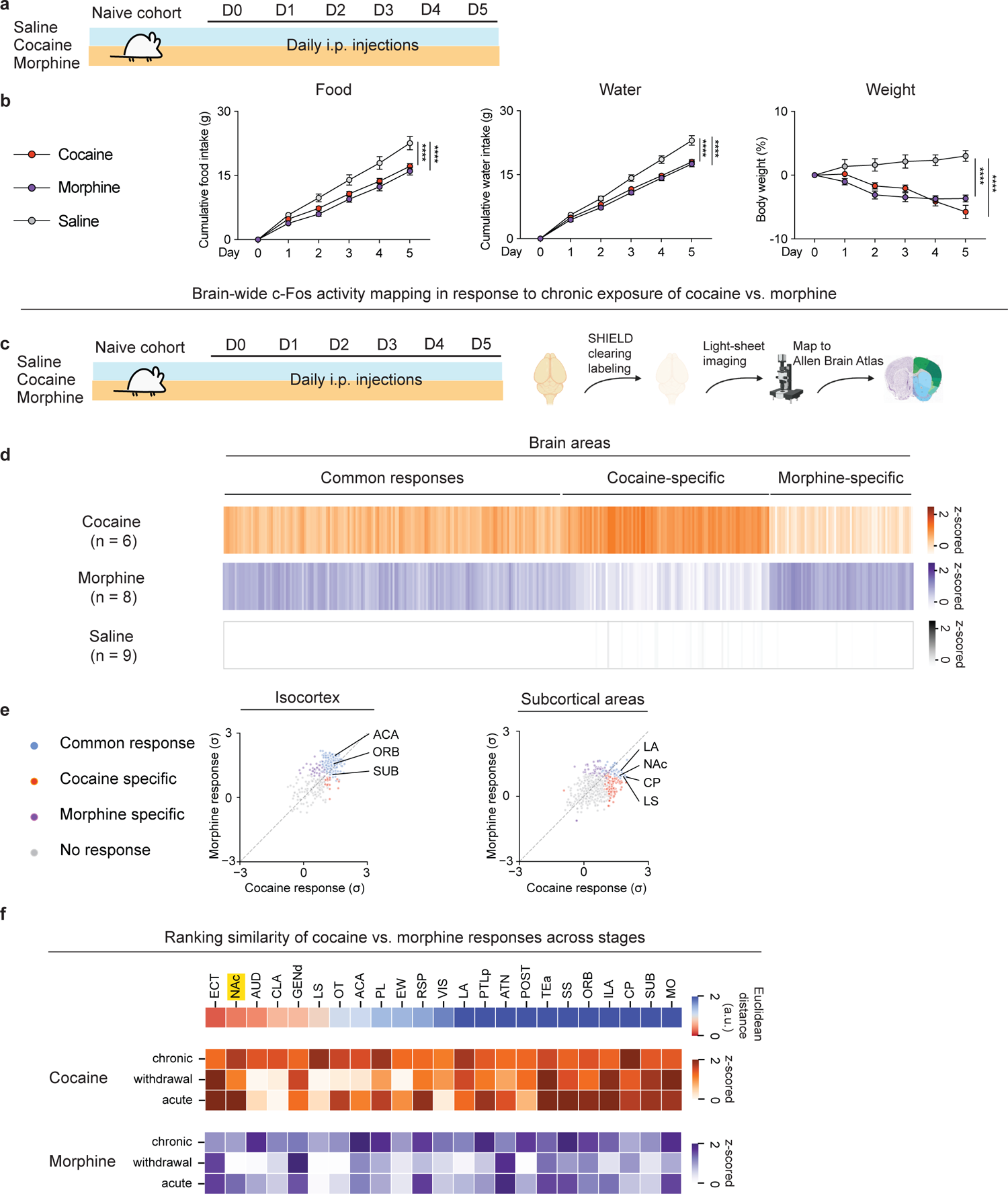
Whole-brain FOS mapping identifies shared regulation of NAc by chronic exposure to cocaine vs. morphine. (a), Schematic of the experimental design for repeated exposure of drug rewards vs. saline. Three cohorts of mice received 20 mg/kg cocaine, 10 mg/kg morphine, saline, respectively via i.p. injections for 5 days. All cohorts had adlibitum access to food and water. Comparisons of (b), Cumulative food intake (g), Cumulative water intake (g), Weight (%) over the 5-day treatment (n = 10, 10, 10 for saline, cocaine, morphine group, respectively, two-way ANOVA with Dunnett’s multiple comparisons). (c), Schematic of the experiment design for chronic exposure of drug rewards vs. saline followed by whole-brain clearing and mapping to Allen Brain Atlas. Three cohorts of mice received saline, 20 mg/kg cocaine, 10 mg/kg morphine, respectively (n = 9, 6, 8 for each group) via i.p. injections for 5 days. (d), Heatmap overview of brain areas showing significant FOS activity across three groups (One-way ANOVA for each brain area with cut-off p < 0.05 classified as statistically significant, followed by K-means clustering). (e), Scatter plot of FOS activity of cortical areas in response to cocaine vs. morphine (left). Scatter plot of FOS activity of subcortical areas in response to cocaine vs. morphine (right). Common response: areas showed significant changes (P < 0.05) of FOS+ counts in cocaine and morphine groups compared to the saline group; Cocaine or Morphine specific: areas only showed significant changes of FOS+ counts in either the cocaine or morphine group compared to the saline group. (f), Similarity of FOS responses across different phases of exposure to cocaine vs. morphine (top). Heatmap representations of brain areas after acute and chronic exposure to cocaine or morphine and after spontaneous withdrawal (bottom). All error bars represent mean ± s.e.m. NS, not significant, *P < 0.05, **P < 0.01, ***P < 0.001, ****P < 0.0001.

Having shown that drug treatment and withdrawal alter food and water intake, we next identified brain regions that could serve as a site that integrates neural responses to natural and drug rewards. To identify brain areas that exhibit convergent and divergent responses to chronic cocaine or morphine exposure, we employed brain-wide FOS mapping after acute administration of either drug (Fig. 1c). Using a SHIELD-based whole-brain clearing approach (Fig. 1c), we observed three general patterns of FOS activation: regions showing cocaine-specific responses, regions showing morphine-specific responses, and regions showing common responses (Fig. 1d). Our goal was to identify brain areas that are activated by both drugs of abuse that might suppress consumption of natural rewards. Cortical regions activated by both drugs included three canonical addiction-related regions: anterior cingulate area (ACA), orbital area (ORB; orbitofrontal cortex), and subiculum (SUB). Of the subcortical areas identified, the lateral amygdala nucleus (LA), nucleus accumbens (NAc), caudoputamen (CP), and lateral septum (LS) also showed common responses to both drugs (Fig. 1e). To identify brain regions showing an augmented response after multiple doses of morphine or cocaine, we performed additional section-based brain-wide FOS mapping experiments to compare activity across distinct phases of drug exposure: acute, chronic, and after spontaneous withdrawal (Fig. S1e, f). Clustering analysis identified several brain areas showing increased FOS activity corresponding to each phase, with a subset of brain areas showing greater activation after acute vs. chronic drug treatment (Fig. S1e). These included the NAc, LA, claustrum (CLA), and others (Fig. 1f). This necessitated pair-wise comparisons of FOS activity from brain areas that show similar responses to both drugs across phases of drug exposure. We found that, among hundreds of brain areas analyzed, the NAc primarily showed increased FOS activity in both phases of chronic and acute exposure to cocaine or morphine (Fig. 1f). Moreover, NAc was one of the top-ranked brain regions exhibiting analogous patterns of responses to cocaine vs. morphine across distinct phases. While consistent with previous findings showing the NAc as a central hub for both cocaine and morphine action (*11–13*), this unbiased, brain-wide analysis further positions the NAc as one of the most prominent convergent targets of drug action in the whole brain.

To link drug action within the NAc to drug influences on homeostatic need, we next tested the effects of gain- or loss-of-function NAc on food and water intake. The NAc houses two key subregions, the core and shell, which exhibit unique cell-type composition, circuit projections, and functions in motivation and learning (*3*). Here, we specifically focus on the NAc core subregion based on its well-established role in coordinating motivated behavior and our prior report demonstrating convergent representations of hunger and thirst in this subregion (*7, 14*). We stereotaxically delivered an inhibitory DREADD, AAV5-hsyn-hM4Di-mCherry, or a control AAV5-hsyn-mCherry bilaterally to the NAc, and three weeks later assessed food and water intake after combined treatment with clozapine N-oxide (CNO) plus cocaine, morphine, or saline (Fig. S2a). We found that chemogenetic silencing of NAc neurons prevented the cocaine- and morphine-induced reductions in food intake, water intake, and body weight during chronic drug exposure (Fig. S2b). This chemogenetic silencing also blocked the suppressive effects of the drugs of abuse on refeeding and rehydration during spontaneous withdrawal (Fig. S2c, d). In contrast, silencing NAc neurons in the control group that received saline without drug exposure had no effect on food or water intake (Fig. S2e).

The principal projection neurons within the NAc are medium spiny neurons (MSNs) which predominantly express either D1 or D2 dopamine receptors. These D1 and D2 populations exhibit a distinct input-output architecture and are known to be differentially engaged by drugs of abuse and natural rewards (*15–18*). Because cocaine and morphine increased FOS in the NAc, we dissected the functional role of these two cell types in coordinating interactions between drug and natural rewards using an optogenetic approach in behavioral paradigms measuring key motivational outcomes. We first stereotaxically delivered AAV5-hsyn-FLEX-ChR2 bilaterally in NAc of D1-Cre or D2-Cre transgenic mice, followed by implantations of optic fibers above the injection sites. We then optogenetically activated D1 or D2 MSNs in fasted mice with free access to food for 10 min, followed by 10 min laser off (Fig. S3a). Activation of D1 or of D2 MSNs potently decreased food intake in fasted mice during refeeding. Additionally, mice with prior activation of D2 MSNs showed a compensatory overeating after laser stimulation, while this was not observed in mice with prior activation of D1 MSNs (Fig. S3a). Similarly, acute activation of D1 or D2 MSNs potently decreased water intake in dehydrated mice (Fig. S3b). In this case, we observed rebound water consumption after laser termination in both D1 and D2 mice (Fig. S3b).

We then tested the influence of D1 or D2 MSN activation on generalized locomotor activity. Activation of D1 neurons caused substantially elevated locomotor activity in *ad libitum* fed mice, while activation of D2 neurons substantially decreased locomotor activity (Fig. S3c), consistent with prior reports (*18–21*). We also tested whether activation of D1 or D2 neurons conveyed valence signals independent of overall locomotion using real-time-place-preference (RTPP). Consistent with previous reports (*18*), we found that activation of D1 neurons increased preference of animals for the stimulation side, indicative of D1 neurons conferring positive valence, while activation of D2 neurons caused an avoidance response to the stimulation side, indicative of conferring negative valence (Fig. S3d).

### Drugs and natural rewards activate an overlapping set of NAc neurons

Thus far, our results raised the possibility that drugs of abuse might alter consumption of natural rewards by modulating the activity of NAc D1 and D2 neurons. To determine whether drugs and natural rewards activate separable or overlapping populations of neurons within the NAc, we recorded the activity of individual neurons in response to food and water deprivation vs. acute or chronic administration of cocaine or morphine. We directly tracked NAc D1 and D2 neuronal activity at single-cell resolution using GRIN lens-based two-photon calcium imaging in headbar-fixed, treadmill-running mice (Fig. S4a, b), as we have recently described (*14*). We designed a series of experiments in which we first recorded individual neuronal responses during feeding and drinking after food or water deprivation, then before and after administration of drugs of abuse for 5 consecutive days, and followed by imaging the same sets of neuronal responses to food and water during drug withdrawal (Summarized in Fig. 2a). In these studies, the response of D1 and D2 neurons to food and water consumption after a period of deprivation were similar to our previous report (*14*).

**Figure. 2.**
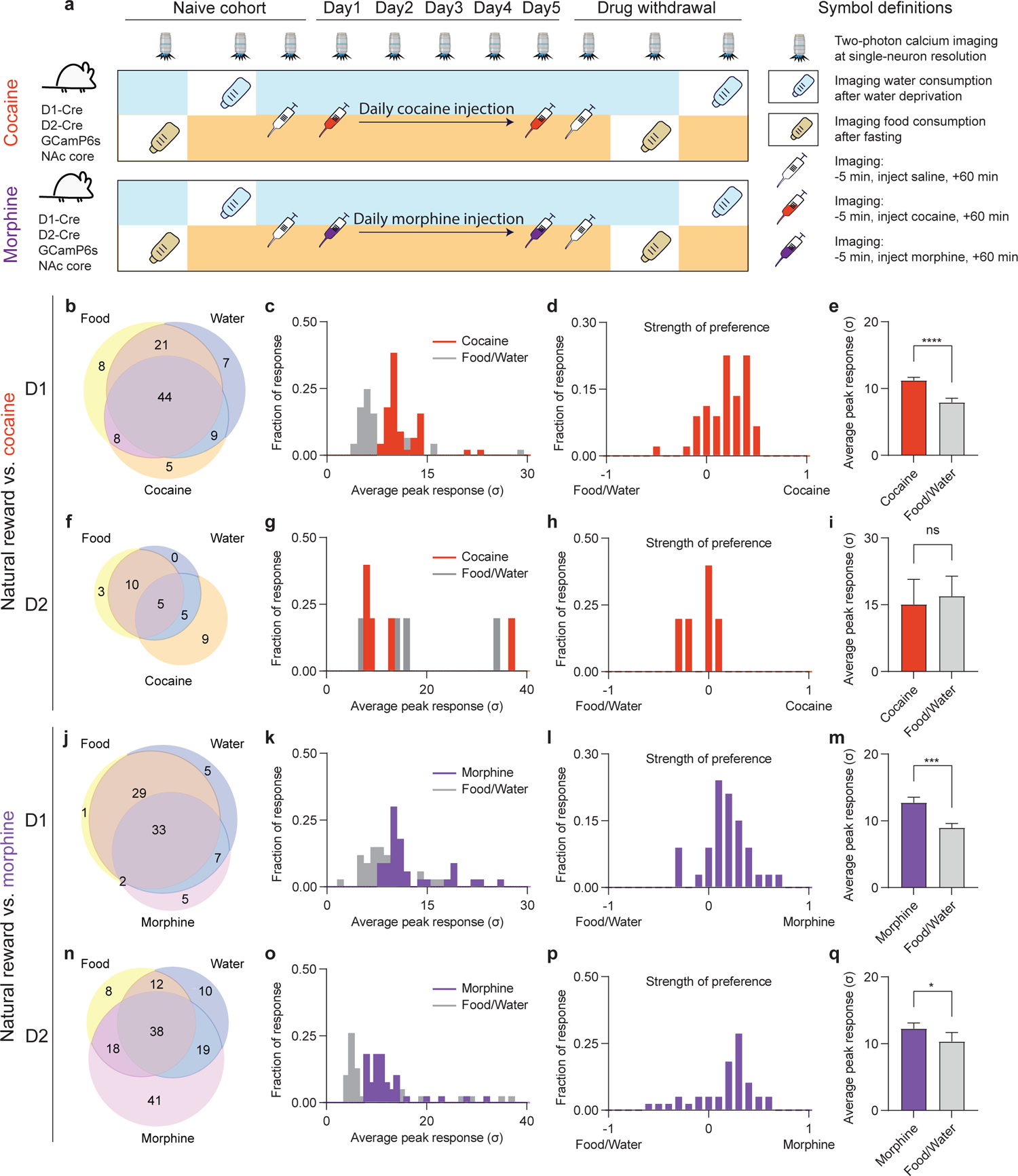
NAc D1 and D2 neurons exhibit common physiological activity across natural and drug rewards. (**a**), Schematic of the experimental design for comparing neuronal responses to natural vs. drug rewards. (b), Venn diagram of activated D1 neurons among food, water, cocaine. N = 111 neurons pooled from 3 mice across all sessions recorded. (c), Histogram distribution of averaged peak responses of the D1 neurons activated by food, water and cocaine. (d), Histogram distribution of preferential activation strength of D1 neurons between food/water vs. cocaine. (e), Comparison of the peaked responses of the above activated neurons (n = 44 neurons, two-tailed Wilcoxon test). (f), Venn diagram of activated D2 neurons among food, water, cocaine. n = 46 neurons pooled from 3 mice across all sessions recorded. (g), Histogram distribution of averaged peak responses of the D2 neurons activated by food, water and cocaine. (h), Histogram distribution of preferential activation strength of D2 neurons between food/water vs. cocaine. (i), Comparison of the peaked responses of the above activated neurons (n = 5 neurons, two-tailed Wilcoxon test). (j), Venn diagram of activated D1 neurons among food, water, morphine. n = 85 neurons pooled from 3 mice across all sessions recorded. (k), Histogram distribution of averaged peak responses of the D1 neurons activated by food, water and morphine. (l), Histogram distribution of preferential activation strength of D1 neurons between food/water vs. morphine. (m), Comparison of the peaked responses of the above activated neurons (n = 33 neurons, two-tailed Wilcoxon test). (n), Venn diagram of activated D2 neurons among food, water, morphine. n = 170 neurons pooled from 3 mice across all sessions recorded. (o), Histogram distribution of averaged peak responses of the D2 neurons activated by food, water and morphine. (p), Histogram distribution of preferential activation strength of D2 neurons between food/water vs. morphine. (q), Comparison of the peaked responses of the above activated neurons (n = 38 neurons, two-tailed Wilcoxon test). All error bars represent mean ± s.e.m. NS, not significant, *P < 0.05, **P < 0.01, ***P < 0.001, ****P < 0.0001.

Cocaine administration elicited a pattern of D1 activation with a high proportion of overlap with those neurons activated by food or water (Fig. 2b). Strikingly, only five out of a total of 111 detected D1 neurons (4.5%) were activated by cocaine alone. In order to precisely compare the magnitudes of neural responses to food and water vs. cocaine, we quantified the Ca^2+^ transient peak amplitudes post food and water consumption vs. post cocaine administration. Among the D1 neurons activated by all three types of rewards (i.e., food, water, and cocaine), the responding neurons were more potently activated by cocaine with highly significant greater peak amplitudes (Fig. 2d, e, cocaine vs. food/water, Peak response: 11.2 ± 0.5, 7.9 ± 0.6, p<0.0001). Cocaine also activated an ensemble of D2 neurons, but a much smaller number compared to D1 neurons. For D2 neurons, the extent of overlap was not significant (Fig. 2f, 5 food-activated and 10 water-activated vs. 19 cocaine-activated D2 neurons) and there was no difference in activation amplitude (Fig. 2h, i).

Similar to cocaine, there was extensive overlap between morphine-activated D1 neurons that responded to food or water (Fig. 2j, 33 food-activated and 40 water-activated vs. 45 morphine-activated D1 neurons) and only five out of a total of 85 D1 neurons (6.2%) were activated solely by morphine. For D2 neurons, the overlap between morphine and natural rewards was partial, with morphine specifically activating 41 neurons vs. 75 neurons that also responded to food or water (Fig. 2n). In contrast to cocaine’s greater activation of D1 neurons than D2 neurons, morphine elicited a roughly equivalent cellular response in D1 and D2 neurons, both of which exceeded responses to natural rewards (Fig. 2k-m, D1 neurons: morphine vs. food/water, Peak response: 12.7 ± 0.8, 9.0 ± 0.7, p<0.001; Fig. 2o-q, D2 neurons: morphine vs. food/water, Peak response: 12.3 ± 0.9, 10.3 ± 1.4, p<0.05). Thus, while cocaine and morphine activate a highly overlapping population of D1 neurons compared to natural rewards, both elicit a significantly greater cellular response. Furthermore, consistent with prior reports (*22*), we show that effects in D2 neurons largely differentiate these psychostimulant and opioid drugs, potentially leading to separable effects within distinct input-output pathways at the circuit level.

In addition to determining the magnitudes of D1 and D2 neuronal responses to cocaine and morphine, we analyzed the correlation of their activity patterns, another key feature of neural dynamics. Because cocaine and morphine both increase locomotor responses in mice and striatal neurons functionally control both locomotor activity and reward processing (*19, 21*), we first segregated out neurons based on whether their activity was associated with non-directed general locomotor activity. To do this, we compared the level of motor activity while a head-fixed mouse was on a treadmill to the neural responses in the cocaine- or morphine-responsive ensembles as either motor-associated or nonmotor-associated neurons, the latter being candidates for neurons that are involved in reward-processing (Fig. S4c, d). We made this assignment in a non-biased manner, but in retrospect noted clear anatomic differences in the focal plane between the motor- and nonmotor-associated ensembles (Fig. S4e), suggesting that distinct motor- and putative reward-related local circuits may be segregated within the NAc. We then evaluated how the nonmotor neurons are modulated by cocaine and morphine by quantifying the correlation in firing in a series of pair-wise comparisons of the activity of individual neurons at each session. We observed that cocaine selectively synchronized (i.e., increased the activity correlation between) nonmotor-associated D1 neurons (Fig. S4f. Connectivity index (cocaine vs. saline): 20 min, 1.54 ± 0.05, 0.86 ± 0.01, p<0.01; 30 min, 1.46 ± 0.14, 0.97 ± 0.11, p=0.054; 40 min, 1.46 ± 0.26, 0.84 ± 0.06, p<0.05), but did not elicit a significant effect on motor-associated neurons (Fig. S4h). By contrast, cocaine did not synchronize the activity of D2 neurons (Fig. S4f). On the other hand, morphine selectively synchronized the nonmotor D2 neurons (Fig. S4g, Connectivity index (morphine vs. saline): 5 min, 1.07 ± 0.22, 0.83 ± 0.07, p<0.05; 10 min, 1.36 ± 0.23, 0.75 ± 0.04, p<0.05; 40 min, 1.32 ± 0.29, 0.75 ± 0.07, p<0.05; 50 min, 1.46 ± 0.21, 0.68 ± 0.09, p<0.01), but did not show a synchronizing effects on D1 neurons (Fig. S4g, i). In aggregate, these data show that neuronal networks that engage in reward processing are anatomically and spatially segregated in NAc from neurons that are associated with motor output. Furthermore, cocaine increases synchronous firing of nonmotor-associated D1 neurons, while morphine progressively induces synchrony of nonmotor-associated D2 neurons, revealing distinct cell-specific responses to the two different drugs of abuse.

### Repeated drug exposure tunes cell-type-specific NAc dynamics

Repeated administration of drugs of abuse causes dynamic, cumulative plasticity within the NAc that has been suggested to cause the behavioral abnormalities underlying addiction (*1, 8, 12, 23, 24*). To examine whether chronic exposure to cocaine or morphine causes neuroplastic changes within ensembles of D1 and D2 neurons, we utilized tensor component analysis (TCA, Fig. 3a), which reduces and organizes multi-trial neuronal dynamics into lower-dimensional factors, effectively subgrouping the responding neurons based on their within-trial dynamics and across-trial evolution (*25*). This analysis revealed that a subset of D1 neurons showed an amplification of their response to cocaine over five days (Fig. 3b; Neuronal response (trial 1 vs. trial 5): 0.02 ± 0.07, 0.36 ± 0.10, p<0.01), while a separate group of D1 neurons that did not respond to acute cocaine in fact showed decreasing activity over the course of the treatments (Fig. 3c). Chronic exposure to cocaine did not show an amplifying effect on D2 neuron activity (Fig. 3f, g; Neuronal response (trial 1 vs. trial 5): 0.44 ± 0.26, 0.17 ± 0.13, p=0.24). In contrast, chronic morphine exposure amplified subsets of both D1 and D2 neural responses over time (Fig. 3j, n, D1 neuronal response (trial 1 vs. trial 5): 0.27 ± 0.10, 0.77 ± 0.12, p<0.0001; D2 neuronal response (trial1 vs. trial 5): 0.15 ± 0.06, 0.39 ± 0.07, p<0.01), while, similar to cocaine, the inactive D1 neural cluster showed a decreasing trend over trials (Fig. 3k). The distinct augmentation of neural responses induced by chronic exposure to cocaine vs. morphine identifies drug-specific pathological changes in D1 and D2 neuronal subtypes within the NAc. Furthermore, this shift in ensemble dynamics, potentially reflective of real-time plasticity, was not observed throughout serial testing with natural rewards (*14*). Thus, drugs of abuse increasingly modulate ensemble structure and dynamics of D1 and D2 neurons, which is not observed in the case of natural rewards, and indicates a progressive alteration of the activity of the associated basal motivation circuits.

**Figure. 3.**
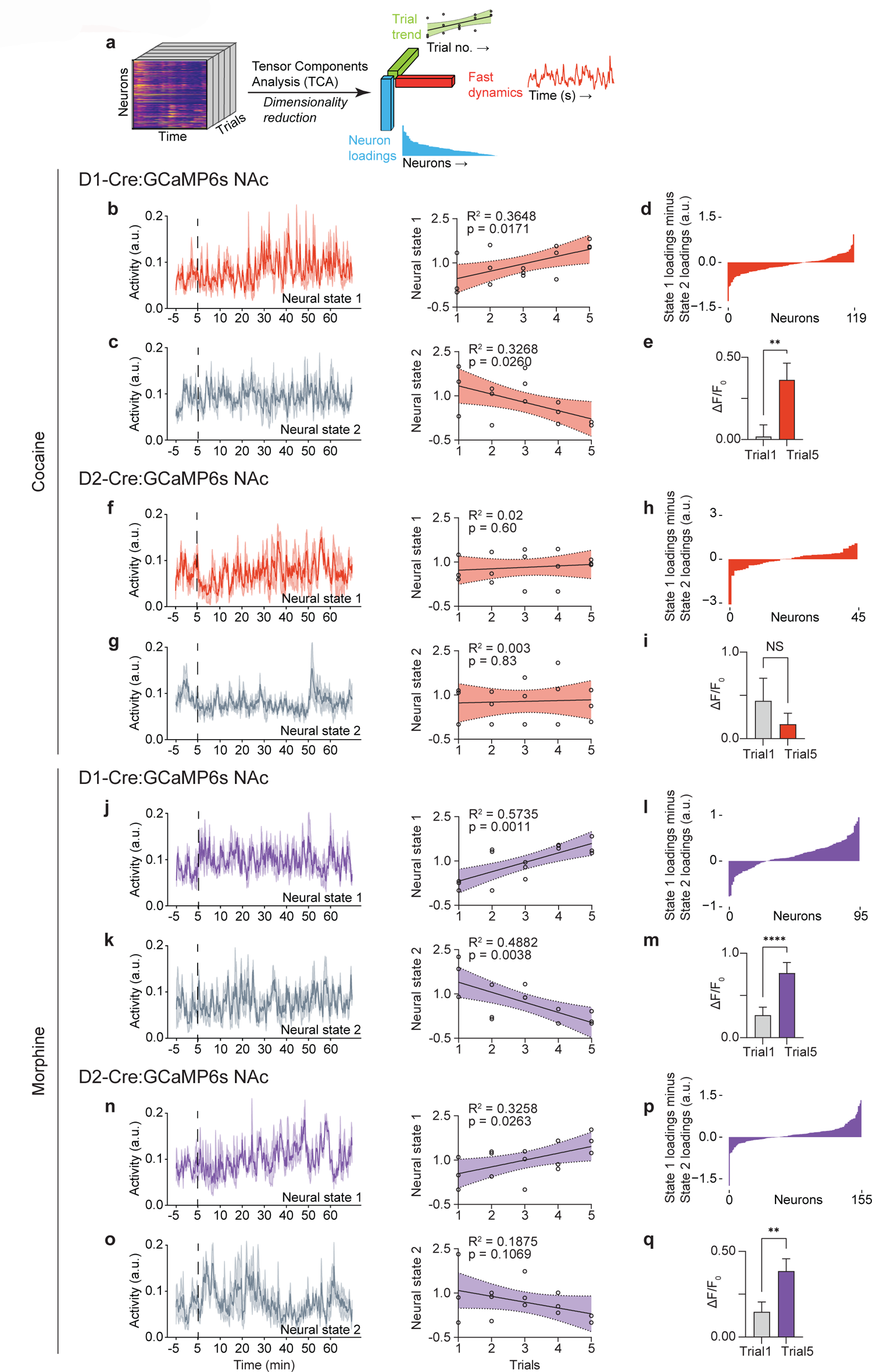
Repeated cocaine or morphine exposure induces and amplifies NAc D1 and D2 neural activity. (**a**), Schematic of tensor component analysis (TCA). (b), Neural state 1 of D1 neurons in response to cocaine and linear regression of the representation of neural state 1 across all sessions. (c), Neural state 2 of D1 neurons in response to cocaine and linear regression of the representation of neural state 2 across all sessions. (d), Loading factors of neurons contributing to state 1 relative to state 2 (n = 119 neurons merged from 3 mice across all 15 sessions). (e), Comparison of D1 neurons positively contributing to neural state 1 between session 1 (day 1) and session 5 (day 5) in response to cocaine (n = 46 neurons, two-tailed Wilcoxon test). (f), Neural state 1 of D2 neurons in response to cocaine and linear regression of the representation of neural state 1 across all sessions. (g), Neural state 2 of D2 neurons in response to cocaine and linear regression of the representation of neural state 2 across all sessions. (h), Loading factors of neurons contributing to state 1 relative to state 2 (n = 45 neurons merged from 3 mice across all 15 sessions). (i), Comparison of D2 neurons positively contributing to neural state 1 between session 1 (day 1) and session 5 (day 5) in response to cocaine (n = 26 neurons, two-tailed Wilcoxon test). (j), Neural state 1 of D1 neurons in response to morphine and linear regression of the representation of neural state 1 across all sessions. (k), Neural state 2 of D1 neurons in response to morphine and linear regression of the representation of neural state 2 across all sessions. (l), Loading factors of neurons contributing to state 1 relative to state 2 (n = 95 neurons merged from 3 mice across all 15 sessions). (m), Comparison of D1 neurons positively contributing to neural state 1 between session 1 (day 1) and session 5 (day 5) in response to cocaine (n = 68 neurons, two-tailed Wilcoxon test). (n), Neural state 1 of D2 neurons in response to morphine and linear regression of the representation of neural state 1 across all sessions. (o), Neural state 2 of D2 neurons in response to morphine and linear regression of the representation of neural state 2 across all sessions. (p), Loading factors of neurons contributing to state 1 relative to state 2 (n = 155 neurons merged from 3 mice across all 15 sessions). (q), Comparison of D2 neurons positively contributing to neural state 1 between session 1 (day 1) and session 5 (day 5) in response to cocaine (n = 97 neurons, two-tailed Wilcoxon test). All error bars represent mean ± s.e.m. NS, not significant, *P < 0.05, **P < 0.01, ***P < 0.001, ****P < 0.0001.

### Withdrawal from drug exposure disorganizes neural representations of natural rewards

A hallmark of addiction is the propensity for relapse after periods of abstinence, suggesting that prior consumption of drugs of abuse causes protracted disruptions in motivation and reward processing. We therefore examined whether withdrawal from repeated drug exposure interferes with neural representations of homeostatic need for food and water. Two days after the last drug injection (i.e., on day 7), mice were fasted (or dehydrated) and allowed to consume food (or water), while the same D1 or D2 neurons were imaged. In a previous study, we found that during refeeding three distinct clusters are activated during the consummatory phase correlating with meal initiation, continued feeding, and meal cessation (*14*). Here, we found that during cocaine withdrawal these three D1 neuronal clusters were entirely disordered with significantly reduced variances. This dramatic change was captured by a reduced percentage of activated D1 neurons and a diminution of correlated neuronal activity, with the reduced variance ratio explained by the top three principal components (PCs) representing the structure of neural dynamics (Fig. 4a-e). However, cocaine withdrawal did not alter D2 neuronal responses to natural rewards (Fig. 4f-j). In contrast, during acute morphine withdrawal, D2 neurons showed markedly increased activation with increased variance ratio explained by the top three PCs (Fig. 4p-t), while morphine withdrawal did not alter the responses of D1 neurons (Fig. 4k-o). Thus, cocaine withdrawal is associated with hypoactivation of the D1 response to natural rewards, whereas morphine withdrawal is associated with hyperactivation of the D2 response to natural rewards. Considering our data above (Fig. S3d, and our data and prior reports showing that D1 neurons promote positive valence and D2 neurons promote negative valence (*18*), these results offer an explanation of separable neural mechanisms of withdrawal across psychostimulants and opioids to produce a negative-valence state.

**Figure. 4.**
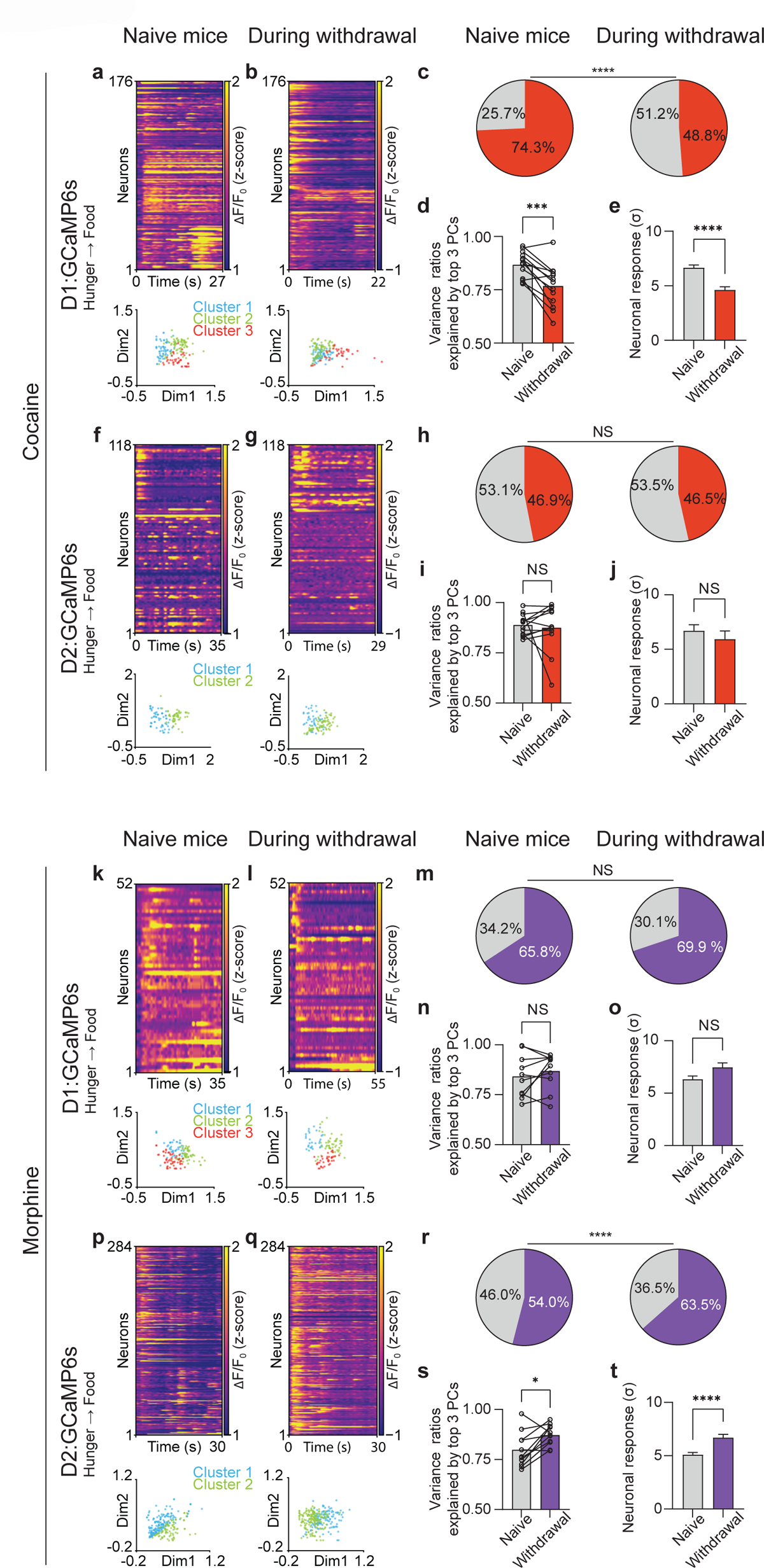
Preferential, cell-type-specific alteration of cocaine- or morphine-induced neuronal responses to natural reward. (**a**), Top: Representative heatmap of D1 neuronal responses during food consumption from an example mouse before exposure to cocaine (n = 176 matched neurons). Bottom: non-negative matrix factorization (NMF) representation of neuronal states labeled by clusters. (b), Top: Representative heatmap of D1 neuronal responses during food consumption from an example mouse during acute withdrawal of cocaine (n = 176 matched neurons). Bottom: NMF representation of neuronal states labeled by k-means clusters. (c), Percentage of D1 neurons activated by food and water consumption in the same mice before and after cocaine exposure (n = 914 matched neurons combined from 13 matched sessions from 3 mice, Fisher’s exact test). (d), Variances explained by top 3 principal components (PCs) during food and water consumption in the same mice before and after cocaine exposure (n = 13 matched sessions from 3 mice). (e), Matched D1 neuronal responses in the same mice before and after cocaine exposure (n = 914 matched neurons, two-tailed Wilcoxon test). (f), Top: Representative heatmap of D2 neuronal responses during food consumption from an example mouse before exposure to cocaine (n = 118 matched neurons). Bottom: NMF representation of neuronal states labeled by clusters. (g), Top: Representative heatmap of D2 neuronal responses during food consumption from an example mouse during acute withdrawal of cocaine (n = 118 matched neurons). Bottom: NMF representation of neuronal states labeled by clusters. (h), Percentage of D2 neurons activated by food and water consumption in the same mice before and after cocaine exposure (n = 488 matched neurons combined from 12 sessions from 3 mice, Fisher’s exact test). (i), Variances explained by top 3 PCs during food and water consumption in the same mice before and after cocaine exposure (n = 12 matched sessions from 3 mice). (j), Matched D2 neuronal responses in the same mice before and after cocaine exposure (n = 488 matched neurons, two-tailed Wilcoxon test). (k), Top: Representative heatmap of D1 neuronal responses during food consumption from an example mouse before exposure to morphine (n = 52 matched neurons). Bottom: NMF representation of neuronal states labeled by clusters. (l), Top: Representative heatmap of D1 neuronal responses during food consumption from an example mouse during acute withdrawal of morphine (n = 52 matched neurons). Bottom: NMF representation of neuronal states labeled by clusters. (m), Percentage of D1 neurons activated by food and water consumption in the same mice before and after morphine exposure (n = 587 matched neurons combined from 10 sessions from 3 mice, Fisher’s exact test). (n), Variances explained by top 3 PCs during food and water consumption in the same mice before and after morphine exposure (n = 10 matched sessions from 3 mice). (o), Matched D1 neuronal responses in the same mice before and after morphine exposure (n = 587 matched neurons, two-tailed Wilcoxon test). (p), Top: Representative heatmap of D2 neuronal responses during food consumption from an example mouse before exposure to morphine (n = 284 matched neurons). Bottom: NMF representation of neuronal states labeled by clusters. (q), Top: Representative heatmap of D2 neuronal responses during food consumption from an example mouse during acute withdrawal of morphine (n = 284 matched neurons). Bottom: NMF representation of neuronal states labeled by cluster. (r), Percentage of D2 neurons activated by food and water consumption in the same mice before and after morphine exposure (n = 1174 matched neurons combined from 11 sessions from 3 mice, Fisher’s exact test). (s), Variances explained by top 3 PCs during food and water consumption in the same mice before and after morphine exposure (n = 11 matched sessions from 3 mice). (t), Matched D2 neuronal responses in the same mice before and after morphine exposure (n = 1174 matched neurons, two-tailed Wilcoxon test). All error bars represent mean ± s.e.m. NS, not significant, *P < 0.05, **P < 0.01, ***P < 0.001, ****P < 0.0001.

### *Rheb* mediates drug-induced interference of natural reward processing

Our calcium imaging findings established a model of neural dynamics wherein drugs of abuse modulate the activity of individual neurons within the NAc that normally process physiologic needs. To explore the potential mechanism by which these drugs alter neural dynamics, we developed an *in silico* approach termed FOS-Seq that leverages the aforementioned brain-wide activity mapping data to search for potential molecular substrates essential for the effects of drug rewards on food and water consumption (*26*). In this analysis, we computationally compared the anatomic distribution of FOS to that of individual genes in the Allen Brain Atlas and computed Pearson Correlation Coefficients (PCC) between brain-wide FOS activity vectors and individual *in situ* expression vectors (ISH vectors) for each gene (Fig. 5a and Fig. S5a) (*27*). Consistent with previous transcriptomic profiling by RNA-seq in response to drugs of abuse (*28, 29*), this approach reliably captured canonical marker genes temporally associated with the addicted state (Fig. 5b, c and Fig. S5b, c). Specifically, after chronic exposure to cocaine, we identified a positive correlation between FOS and *Drd1*, *Drd2*, *Drd3*, *Deaf1*, *Fosb,* and *Lcn2* expression (Fig. 5b). Additionally, we found that the *Oprl1* and *Rxra* genes were negatively correlated with FOS, while the *Oprm1* gene was not correlated with FOS (Fig. 5b). In contrast, chronic exposure to morphine elicited positive correlations between FOS and *Fosb* and *Crebbp* (Fig. 5c). We also found a negative correlation between FOS and *Agt*, *Htr2c*, *Oprl1*, *Oprm1,* and *Oprk1* expression (Fig. 5c). To validate this *in silico* approach, we conducted the same analysis for FOS activity vectors from acute exposure and spontaneous withdrawal of cocaine or morphine. Again, this approach consistently captured canonical marker genes known to show changes in gene expression in response to each of the two drugs of abuse (Fig. S5d, e). To directly identify genes from FOS-Seq that are shared between chronic cocaine and morphine exposure, we generated a scatter plot of PCCs for each gene by analyzing the intersection among those that were significantly correlated with FOS (Fig. 5d). We found that *Rheb* was a top-ranked gene positively correlated with FOS, and uniquely shared by chronic exposure to cocaine or morphine (Fig. 5d). *Rheb* encodes a GTP-binding protein that phosphorylates mTOR and activates downstream pathways (*30*). Consistent with prior reports (*31–36*), we also observed several other genes in the mTOR pathway that are significantly correlated with FOS activity (Fig. S5c, d).

**Figure. 5.**
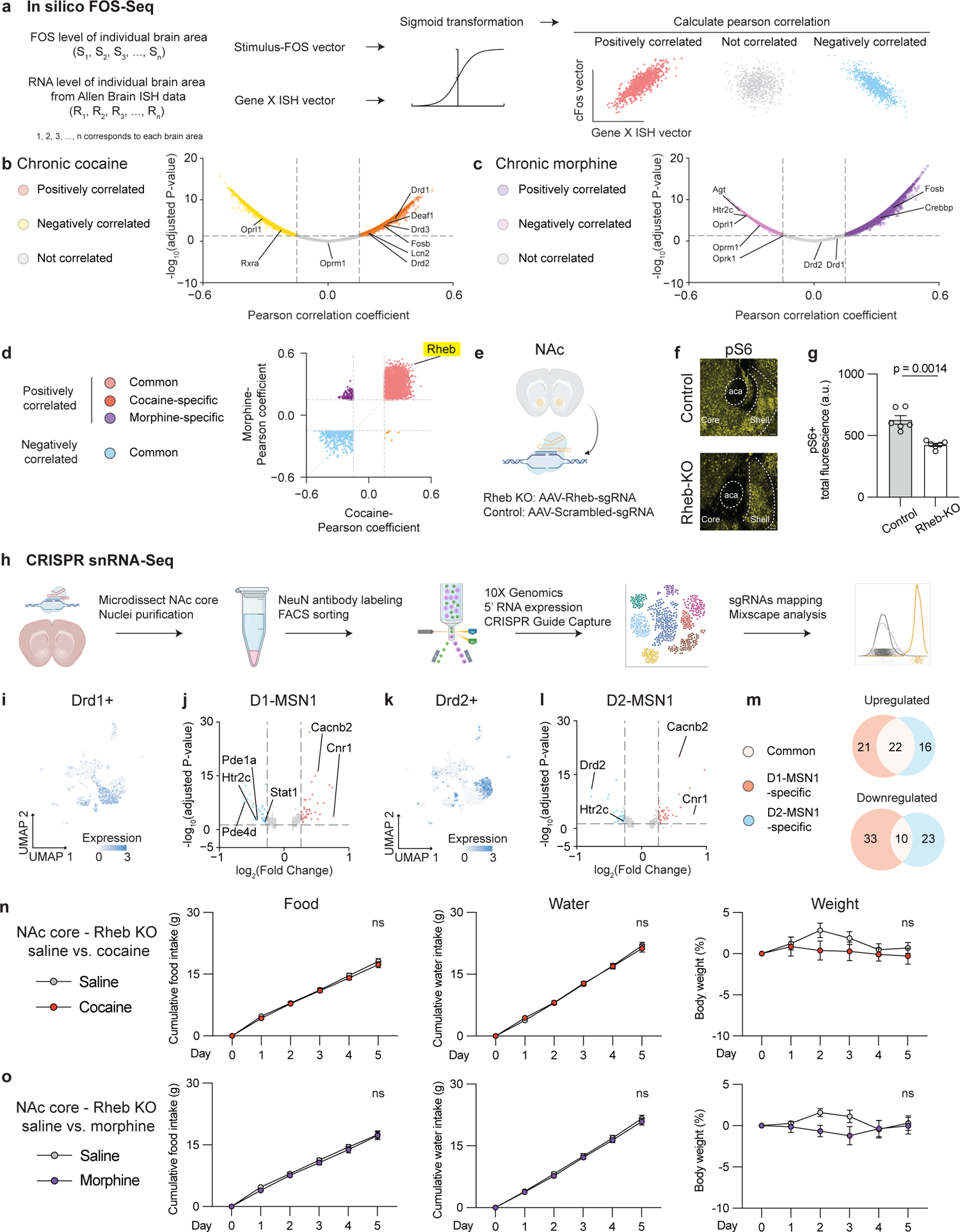
*Rheb* serves as a molecular link to enable disruption of natural reward processing induced by repeated exposure to either cocaine or morphine. (a), Schematic of the *in silico* FOS-Seq approach to identify genes associated with brain-wide c-Fos activity patterns. Genes with Pearson Correlation Coefficient > 0.15 or < −0.15 and p < 0.05 are classified as positively correlated or negatively correlated genes, while the rest are considered to be not correlated. Adjusted P-values are FDR-corrected at 5% threshold. (b), Volcano plot of genes associated with chronic exposure of cocaine. (c), Volcano plot of genes associated with chronic exposure of morphine. (d), Scatter plot of Pearson coefficient from genes associated with chronic exposure of cocaine vs. Pearson coefficient from genes associated with chronic exposure of morphine. (e), Schematic of *in vivo* NAc region-specific knockout of *Rheb* gene by co-expressing Cre and Rheb-sgRNAs or their control scrambled-sgRNAs in NAc core in LSL-Cas9 transgenic mice. (f), Immunohistochemistry validation of pS6 levels in Rheb knockout (KO) vs. Control (WT) at baseline. (g), Quantification of total pS6 fluorescent intensity in the NAc. (h), Schematic of snRNA-seq after CRISPR perturbations (n = 7001 cells mapped with either sgRNAs). (i), Distribution of *Drd1+* cells in the UMAP. (j), Differentially expressed genes in Rheb-perturbed cells vs. control cells in the D1-MSN1 cluster (n = 598, 390 cells respectively). (k), Distribution of *Drd2+* cells in the UMAP. (l), Differentially expressed genes in Rheb-perturbed cells vs. control cells in the D2-MSN1 cluster (n = 449, 375 cells respectively). (m), Venn diagram of genes significantly regulated by *Rheb* perturbation between D1-MSN1 and D2-MSN1 clusters. (n), Comparisons of cumulative food intake (g), water intake (g), weight (%) in the Rheb-KO group treated with saline for 5 days followed by another 5-day cocaine treatment (n = 8, 8 for each group, two-way ANOVA w’th Šídák’s multiple comparisons). (o), Comparisons of cumulative food intake (g), water intake (g), weight (%) in the Rheb-KO group treated with saline for 5 days followed by another 5-day morphine treatment (n = 8, 8 for each group, two-way ANOVA w’th Šídák’s multiple comparisons). All error bars represent mean ± s.e.m. NS, not significant, *P < 0.05, **P < 0.01, ***P < 0.001, ****P < 0.0001.

Considering our finding that inhibition of NAc neurons blunts the ability of drugs of abuse to suppress natural reward consumption (Fig. S2), we next tested whether *Rheb* may have a role in coordinating this effect. To do this, we generated a region-specific knockout of *Rheb* and its control by stereotaxically delivering AAV5-hsyn-Cre and AAV5-Rheb-sgRNAs-hsyn-mCherry or AAV5-control-sgRNAs-hsyn-mCherry bilaterally in NAc core of LSL-Cas9 transgenic mice (Fig. 5e). Using this approach, we first immunohistochemically probed levels of phospho-S6 (pS6; a ribosomal protein), a well-established marker for RHEB-mTOR activity (*37*), in the NAc of the *Rheb* knockout and control mice (Fig. 5f). We found that the *Rheb* knockout (Rheb-KO) group showed decreased pS6 fluorescence in the NAc (core) compared to the control group (Fig. 5f, g), suggesting a high knockout efficiency of *Rheb* by CRISPR *in vivo*. We also conducted single-nucleus RNA-sequencing (snRNA-seq) from neurons in the NAc core and mapped those transduced with sgRNAs post hoc (Fig. 5h). This approach allowed us to examine cell-type-specific perturbation effects of *Rheb* knockout. Consistent with previous studies, our snRNA-seq of NAc core revealed 18 different clusters that were marked by single or double molecular markers illustrated by UMAP and violin plots (Fig. S6a, b). As expected, D1 (*Drd1+*) and D2 (*Drd2+*) cells were spatially segregated in the UMAP (Fig. 5i, k). Among these cells, D1 MSNs were further comprised of three subtypes: cluster1 *Drd1/Pdyn* (D1-MSN1), cluster5 *Drd1/Tshz1* (D1-MSN2), and cluster11 *Drd1/Drd3* (D1-MSN3); D2 MSNs were also comprised of three subtypes: cluster2 *Drd2/P2ry1* (D2-MSN1), cluster7 *Drd2/Reln* (D2-MSN2), and cluster14 *Drd2/Htr7* (D2-MSN3).

Using our snRNA-seq results, we evaluated how CRISPR perturbation affected either D1 or D2 MSNs. Prior reports show that *in vivo* CRISPR perturbation induces both frame-shift and in-frame mutations, thus resulting in subsets of cells that escape perturbations (*38–40*). To parse these ‘escaped’ cells, we applied Mixscape analysis (*38*), a validated algorithm that classifies *Rheb*-sgRNAs+ cells into perturbed and escaped cells compared to Control-sgRNAs+ cells (Fig. S6c). This analysis further revealed that perturbed cells were specific within D1-MSN1 and D2-MSN1 clusters, two main subtypes of D1 and D2 MSNs (Fig. S6d). We next performed differential gene expression analysis on perturbed vs. control cells from D1-MSN1 and D2-MSN1 clusters (Fig. 5j, l). These analyses revealed that *Rheb* perturbation significantly decreased the expression of *Htr2c*, which encodes the 5HT2C serotonin receptor, and increased the expression of *Cacnb2*, which encodes a voltage-gated calcium channel subunit, as well as *Cnr1*, which encodes the CB1 cannabinoid receptor, in both D1-MSN1 and D2-MSN1 clusters (Fig. 5j, l). In contrast to this common set of genes, *Rheb* perturbation in D1-MSN1 specifically decreased *Pde1a* and *Pde4d*, which encode phosphodiesterase enzymes associated with cAMP signal transduction. In D2-MSN1, *Rheb* perturbation specifically decreased the expression of *Drd2* (Fig. 5j, l). Overall, *Rheb* perturbation in D1-MSN1 and D2-MSN1 led to transcriptional regulation of both common and distinct subsets of genes (Fig. 5m). Gene ontology analysis further revealed that distinct pathways within each cell type were affected by *Rheb* perturbation (Fig. S6e). These analyses indicate that *Rheb* regulates cell-type-specific signaling pathways by which drugs of abuse cause neuroplasticity and associated behavioral responses.

Considering that *Rheb* perturbation produces transcriptomic consequences that are associated with drug signaling within the NAc core, we examined whether *Rheb* has a causal role in mediating the behavioral effects of drugs of abuse on natural reward consumption. Mice received viral transduction of Rheb-sgRNAs or control-sgRNAs in NAc core, and were subsequently administered saline for 5 days, followed by an additional 5 days of repeated exposure to cocaine or morphine. We found that, in the knockout group, cocaine and morphine no longer elicited a significant alteration of food intake, water intake, or body weight compared to the prior saline treatment in the same cohort (Fig. 5n, o). In contrast, the control group consistently showed reduced consumption of natural rewards (Fig. S6f, g). These data mirror our prior functional data, and further suggest that increased mTOR signaling caused by repeated exposure to either cocaine or morphine contributes to the augmented cellular responses to drugs of abuse and diminished responses to food or water.

### Identification of NAc input relays for integration of natural and drug reward

The NAc receives inputs from numerous regions that carry information relevant to reward processing across drug classes (*15, 16, 41–43*). In contrast, there are few direct inputs from brain regions that sense physiological needs such as arcuate nucleus neurons that regulate feeding or subfornical organ neurons that regulate fluid intake (*44–46*). Since our data show that NAc neurons are activated by thirst and hunger, we set out to identify how signals reflecting physiological needs input to the NAc. Toward this end, we delivered rAAV2-retro-CAG-GFP virus unilaterally to the NAc to label monosynaptic inputs. Mice were then treated with chronic injections of cocaine, morphine, or saline (Fig. S7 and Fig. S8), followed by SHIELD-based whole-brain mapping of FOS and GFP expression in the same brain. This enabled us to overlay sites of FOS activation with sites that project monosynaptically to the NAc and express endogenous GFP from the retrograde NAc tracer. We then overlaid GFP density and FOS activity in the same brain in response to each stimulus (Fig. S7 and Fig. S8). We found that lateral septum (LS), subiculum (SUB), anterior dorsal thalamus (ATN), prefrontal cortex (ORB, ILA, PL, and ACA), ectorhinal area (ECT), and amygdala (LA, CLA) showed FOS after both cocaine and morphine treatment while also showing strong retrograding GFP signals from NAc (Fig. S9a-c). Intriguingly, the amygdala is known to regulate feeding and drinking, while cortical regions have been suggested to exert higher order control over drug-seeking and drug-taking (*47–50*). To examine the causal roles of these nodes, we conducted gain-of-function chemogenetic activation experiments. We delivered rgAAV-hSyn-DIO-hM3D(Gq) into the NAc core, and delivered AAV5-hSyn-Cre to different mouse groups in these areas: LS, ORB, ventral SUB (vSUB), ATN, ACA, LA, medial PFC (mPFC: ILA, PL), or CLA/ECT (Fig. S9d). This combined retrograde-DREADD and Cre-labeling strategy enables specific expression of an activating-DREADD in NAc-projecting neurons in each of these areas. Subsequently, CNO was administered to each group of mice, and food or water consumption was measured. We found that activation of LS neurons or ORB neurons among these 8 NAc-projecting nodes significantly decreased food consumption. Moreover, only the activation of ORB neurons significantly decreased water consumption (Fig. S9d). Thus, these data support the view that the NAc is a convergent node in a common reward pathway carrying higher-order information relevant to homeostatic value, which drugs of abuse corrupt in a circuit-dependent manner.

## Discussion

Here, we identify a reward scaling system contained within the NAc that normally responds to food and water deprivation and that becomes corrupted by drugs of abuse. Our results implicate crucial molecular and circuit mechanisms that support this process, potentially underlying behavioral dysfunctions associated with drug addiction and withdrawal. We found that two classes of drugs of abuse (cocaine and morphine) and natural rewards not only activate NAc neurons, but engage largely overlapping sets of neurons within this region. This discovery raises a central question: How does the NAc differentiate between natural and drug rewards? A key advantage of two-photon calcium imaging is the ability to deconstruct individual neuron dynamics while tracking the same neuron over days (*14*). Using this approach, we found that NAc neurons distinguish between natural and drug rewards in two ways: the strength of activation, and firing rates over repeated exposure. Compared to the structured, stereotypic neural response in NAc during feeding and drinking (*14*), drug rewards activate a highly overlapping set of neurons but to a greater degree, while amplifying these neuronal responses over repeated exposure and ultimately dysregulating their responses to food and water. This is consistent with our FOS mapping results which, along with the calcium imaging observations, also reveal distinct responses of D1 vs. D2 MSNs over time. These results thus provide a framework within which to understand the basis for the drug-induced neuroplasticity induced in these particular subsets of NAc neurons by repeated exposure to drugs of abuse.

### Divergent cellular dynamics link drug and natural reward representations

In addition to the overlapping sets of neurons that respond to both natural and drug rewards, we also observed unique changes of cellular dynamics in response to cocaine vs. morphine, which differentiated their preferential actions across D1 and D2 NAc neurons.

Consistent with *ex vivo* and histological evidence from previous studies (*22, 51*), we observed that cocaine almost exclusively activates D1 neurons and amplifies their activity with repeated exposure, while morphine activates and amplifies both D1 and D2 neuron function. This cell-type-specific evolution of NAc dynamics with repeated drug exposure could reflect divergent neural substrates that coordinate separable responses across drug classes and that are computed in the same brain region to enhance the reward value of each drug over the value of satisfying physiologic needs. In addition to physiologic symptoms, withdrawal from drug intake produces dysphoria and general negative affect, which is potentially reflective of a rebound from positive to negative valence. We and others have shown that D2 activation can promote aversive effects, while D1 activation promotes positive reinforcement (*17, 18, 52*). Therefore, our finding of diminished D1 neuronal responses to natural rewards during withdrawal from either cocaine or morphine may reflect an interference with appetitive behavior, while augmented D2 neuronal responses during morphine withdrawal may reflect a shift toward negative affect-based seeking. Thus, our results point to partly separable neuronal coding principles across drug classes with direct relevance to addiction and withdrawal.

These physiology results demonstrate a unique ability of drugs of abuse to induce types of neuroplasticity that are not observed for food or water. We found that cocaine and morphine caused clear qualitative and quantitative differences in neuronal activation and tuning with repeated exposure in contrast to both food and water (*14*). Importantly, our studies examine these physiologic responses in food or water deprived states, further implying that drugs of abuse activate these systems to a greater degree than homeostatic need-based signaling. The fact that these changes are not observed for food or water suggests that drug use dysregulates a highly coordinated system that normally matches physiological need to behavior. This dysregulation is evident in our studies showing an effect of morphine and cocaine to reduce food and water intake with distinct effects on NAc neural populations. Similarly, drug withdrawal decreases D1 and increases D2 activity, both of which would contribute to a negative affective state that is no longer properly regulated by natural rewards. This effect could then drive drug-seeking at the expense of natural reward. Overall, these data imply a strong distinction between the rewarding properties of drugs of abuse and natural rewards like food, which have been suggested to be addictive themselves (*9, 53*). Together, these results enforce the idea that food and water intake is a highly regulated homeostatic process while drug addiction is progressive and support the conclusion that cocaine and morphine disrupt and subvert a normal physiologic drive through partly shared and partly distinct neural actions.

### *Rheb* links drug exposure with natural reward processing

Cell-specific calcium activity also drives molecular changes in NAc and, consistent with this, numerous studies have implicated cell-type-specific molecular processes within the NAc that mediate the addictive actions of drugs of abuse (*54*). One example is ΔFOSB, whose D1 MSN-specific induction by chronic cocaine and combined D1 and D2 MSN induction by chronic opioids (*55*) mirrors the patterns of NAc neuronal activation observed here. Interestingly, ΔFOSB is also induced by high levels of consumption of natural rewards and this induction in turn increases such consumption (*56*). However, this mechanism is self-limited for natural rewards, which do not show the amplification of signals over time as do drugs of abuse. This raises the question of whether a specific molecular substrate(s) within the NAc drives the pathological effects of drug exposure while simultaneously interfering with innate responses to natural rewards. Our study indeed identifies such a substrate—the mTOR activator, *Rheb*—as a crucial intermediate specifically linking the ability of drug exposure to attenuate food and water consumption. This finding raises the possibility that sustained mTOR activation in NAc by repeated drug exposure might help orchestrate neural plasticity during the development of addiction. Consistent with this hypothesis, previous studies have also shown that reducing mTOR activity in NAc with rapamycin, a potent mTOR inhibitor, inhibits cue-induced reinstatement of cocaine-seeking without affecting cue-induced sucrose seeking behavior, indicating a specific role for mTOR activity in drug action (*33, 34, 36*). However, these effects appear to be region-specific as reducing mTOR activity in the VTA mediates morphine tolerance (*32*). Our study and prior reports using single-cell sequencing and spatial transcriptomics have revealed considerable cellular heterogeneity within the NAc (*57–61*), but it remains unknown how the same component of the mTOR pathway controls divergent actions of cocaine and morphine. One possibility, implied by our results from imaging and snRNA-seq after CRISPR perturbation, is that these two types of drug rewards preferentially sensitize firing rates of distinct cell types, and that the intracellular mTOR pathway is then essential for each cell-specific drug action. This raises an important future line of research to understand how a generic signal transduction pathway distinguishes between rewarding stimuli and adapts neural sensitivity to each stimulus (*62, 63*). Answering these fundamental questions could potentially lead to development of cell-specific mTOR modulators that would dramatically enhance addiction treatment options.

### Drugs and natural rewards engage similar NAc input architecture

The NAc is an integrative node in the limbic circuitry that is essential for reward processing and generating motivated behavior. These higher order psychological and behavioral processes require integration of complex interoceptive and exteroceptive signals, information relevant to learned associations among sensory stimuli, reward acquisition and anticipatory and coordinated motoric output centers. It is therefore likely that the specific input/output networks connected with the NAc influence the specific information being processed within the NAc. While our results suggest a role for the NAc in sensing hunger and thirst, the NAc receives very few direct ascending inputs from canonical energy and fluid homeostasis nodes such as the hypothalamic arcuate nucleus or subfornical organ, respectively (*45, 64*). We addressed this question by using a whole-brain mapping approach, which provided a comprehensive map of ascending nodes activated by both cocaine and morphine that project to NAc, including prefrontal cortex, LA, anterior thalamic nuclei and LS. Prior literature has implicated several of these regions in both natural and drug reward processing, from homeostatic intake to roles in coordinating conditioned food- or drug-seeking behavior (*47–50, 64, 65*). By activating several of these pathways, we identified a particularly important role for ORB inputs to the NAc in coordinating consumption of natural reward. The ORB is known to be crucial for decision making and updating reward value based on internal states (*66–68*). Thus, our findings suggest that value information communicated to the NAc through the ORB has a broad influence on consummatory behaviors that fulfill homeostatic need. Considering the ability to drugs of abuse to cause persistent changes to ORB function (*69*), it is plausible that this ORB-NAc pathway is crucial for the ability of drugs of abuse to gain access to higher-order information flow that is responsible for coordinating goal-directed behavior. Together, these results point to the NAc as a crucial gateway for integrating and amplifying the value of drug rewards, and suggest that this action serves as a top-down function over behavior towards other goals to ultimately promote drug seeking at the expense of natural, healthy rewards. Future work aimed at further dissecting the NAc circuit architecture that contributes to these effects at the cellular and molecular levels would be informative.

Overall, our findings establish that mechanisms in the NAc that scale intrinsic reward value are overtaken by drugs of abuse and point to a potential molecular substrate supporting this process. These results extend prevailing theories of addiction development and maintenance, and suggest the involvement of partly shared but also partly drug-class-specific mechanisms based on dynamic changes within the NAc at different phases of drug exposure. Importantly, these findings reveal that recurrent exposure to drugs of abuse induces a sustained activation of the RHEB-mTOR pathway leading to disruptions in the cell-type-specific representations of innate needs. Together, these studies provide a framework to understand how drugs of abuse cause neuroplastic changes in brain systems coordinating motivation, ultimately producing behavioral abnormalities relevant to addiction.

## Author contributions

B.T., C.J.B, A.V., J.M.F., and E.J.N conceived the study. B.T. prepared animal specimens and carried out behavioral assays, optogenetic experiments, snRNA-seq sample preparation and analysis. B.T., C.J.B, and T.N. implemented the behavioral assay for functional imaging experiments, performed these experiments and analyzed data. B.T., C.J.B., J.M.F., A.V., and E.J.N. conceived functional imaging aspects of the study, designed and built the imaging system, and advised on data collection and analysis. C.J.B. and E.J.N. assisted with design of drug treatment paradigms and supplied reagents. B.T., C.J.B., T.N., A.V., J.M.F., and E.J.N. wrote the manuscript. All authors discussed the results and contributed to the manuscript. A.V., J.M.F., and E.J.N supervised the project and provided funding support.

## Acknowledgements

We thank Cori Bargmann and Nathaniel Heintz for advice and comments on the manuscript. We thank Estefania Azevedo, Lisa Pomeranz, Sarah Stern, Donghoon Lee, and Qian Lin for comments and advice on behavioral, molecular, imaging experiments, and data analysis. We thank Kristina Hedbacker for illustrations. We thank Inna Piscitello for helping with manuscript submission. We thank Alex Kwan for providing advice and materials on FOS-Seq. We thank James Petrillo at the Precision Instrumentation Facility. We thank Ji-Dung Luo for help with snRNA-seq quality control and preliminary analysis. We thank Helen Duan and Connie Zhao for help with preparing snRNA-seq libraries. We thank the FCRC team for helping with FACS sorting. We thank Zuohang Wu, Chuanyun Xu, and Ke Ding for discussions. B.T. acknowledges support from the David Rockefeller Fellowship. C.J.B. was supported by a postdoctoral fellowship from the National Science and Research Council of Canada. T.N. was supported by a Kavli Fellowship at The Rockefeller University. A.V. acknowledges funding from NINDS (5U01NS115530, 1RF1NS110501, 1RF1NS113251), the National Science Foundation (DBI-1707408), and the Kavli Foundation. J.M.F. acknowledges support from JPB Foundation. E.J.N. acknowledges support from NIDA (P01DA047233, R01DA014133).

## Methods

### Mouse strains

Male adult mice (8-24 weeks old) were used for experiments. We obtained Drd1-Cre mice (Drd1-Cre120Mxu/Mmjax, stock no. 37156), wild-type mice (C57BL/6J, stock no. 000664), and LSL-Cas9 mice (stock no. 026175) from the Jackson Laboratory. Drd2-Cre mice were obtained from Dr. E. Azevedo at The Rockefeller University as previously described (*70*). All mice were maintained in temperature- and humidity-controlled facilities on a 12-h light-dark cycle (light on at 7:00 am) and had ad libitum access to food and water except when noted otherwise. All feeding experiments used standard rodent chow pellets. For fasting or dehydration experiments, mice were overnight fasted or dehydrated (16-24 hours of food or water deprivation). All experimental protocols were approved by the Rockefeller University IACUC, according to the NIH Guide for the Care and Use of Laboratory Animals.

### Viral vectors

The following AAV viruses were purchased from the vector core at the University of North Carolina: AAV5-EF1a-DIO-ChR2-YFP, AAV5-EF1a-DIO-YFP, AAV5-EF1a-DIO-mCherry. The following AAV viruses were purchased from Addgene: AAV5-hsyn-hM4Di-mCherry, AAV5-hsyn-mCherry, rAAV2-retro-CAG-GFP, AAV5-hsyn-Cre, AAV5-AAV1-hSyn-FLEX-GCaMP6s. The plasmid expressing guide RNAs targeting *Rheb*: pAAV-U6-sgRNA1[Rheb]-U6-sgRNA2[Rheb]-hsyn-mCherry, and non-targeted control: pAAV-U6-sgRNA1[Scrambled]-U6-Rheb-sgRNA2[Scrambled]-hsyn-mCherry were designed and engineered at VectorBuilder. The plasmids were packaged into AAV5 at Janelia Viral Tools. sgRNA1[Rheb] sequence: ACCAAGTTGATCACGGTAAA, sgRNA2[Rheb] sequence: GTTCTCTATGGTTGGATCGT. sgRNA1[Scrambled] sequence: GTGTAGTTCGACCATTCGTG, sgRNA2[Scrambled] sequence: GTTCAGGATCACGTTACCGC.

### Stereotaxic surgery

Mice were induced with 3% inhaled isoflurane anesthetic in oxygen, placed in a stereotaxic apparatus (Kopf Instruments) and maintained at 1.5% isoflurane during surgery. The coordinates of injection sites were determined by the website tool: http://labs.gaidi.ca/mouse-brain-atlas. Viruses were bilaterally injected in the nucleus accumbens core (300-500 nl per side, 100 nl/min) using the following coordinates relative to the bregma: AP: +1.32; ML: ±1.1; DV: −4.25-4.5. For optogenetic experiments, optic fibers with 200-μm diameter core (Thorlabs CFML12U-20) were placed 0.3-0.5 mm above the virus injection site. For in vivo two-photon imaging surgeries, in order to obtain sufficient expression of GCaMP6s, 600 nl of virus were delivered (300 nl per coordinate) at AP: +1.22; ML: +1.2; DV: −4.25 and AP: +1.42; ML: +1.2; DV: −4.25 or at AP: +1.32; ML: +1; DV: −4.25 and AP: +1.32; ML: +1.3; DV: −4.25. A gradient-index (GRIN) lens with 1 mm diameter and 4.38 mm length (GRINTECH NEM-100-25-10-860-S) was implanted at the following coordinates: AP: +1.32; ML: +1.2; DV: −4.00, 0.2 mm above the injection site. Mice were used for behavioral experiments 2-3 weeks after surgeries. For in vivo two-photon imaging experiments, mice were used no earlier than four weeks after surgery to allow sufficient and stable GCaMP6s expression. For retrograde activating DREADD experiments, 500 nl of AAV5-hSyn-Cre virus were delivered bilaterally at each NAc-projecting area with following coordinates: LS (AP: 1.0, ML: ±0.4, DV: −3.5), ACA (AP: 1.0, ML: ±0.3, DV: −1.5), CLA/ECT (AP: 1.0, ML: ±2.8, DV: −3.68), mPFC (AP: 2.0, ML: ±0.4, DV: −2.5), ORB (AP: 2.6, ML: ±1.65, DV: −2.8), vSUB (AP: −4.1, ML: ±3.3, DV: −3.8), ATN (AP: −0.75, ML: ±0.9, DV: −3.15).

### Histology

Mice were transcardially perfused with PBS followed by 10% formalin or 4% PFA. Brains were dissected and post-fixed in 10% formalin or 4% PFA at 4°C overnight. Brains were sectioned into 50-μm or 100-μm coronal slices using a vibratome (Leica). For immunohistochemistry, brain sections were blocked (0.1% Triton X-100 in PBS, 3% bovine serum albumin, 2% donkey serum) and then incubated with primary antibody (rabbit anti-c-fos, Cell Signaling, 1:500 for 100-μm sections; rabbit anti-Phospho-S6, Invitrogen, 1:1000 for 50-μm sections) for 2 days at 4°C. Sections were then washed and incubated with secondary antibody (donkey anti-rabbit IgG Alexa 568, Invitrogen, 1:500 for 100-μm sections; donkey anti-rabbit IgG Alexa 647, Invitrogen, 1:1000 for 50-μm sections) for 1 hour at room temperature, washed again, mounted with DAPI Fluoromount-G (Southern Biotech) and imaged with SlideView microscope (VS200, Olympus). Images underwent minimal processing (such as adjusting brightness and contrast) performed using ImageJ.

### Whole-brain FOS activity mapping and GFP labeling

SHIELD-based whole-brain clearing and labeling was employed for mapping FOS activity and GFP density in mice that had received chronic exposure of cocaine, morphine vs. saline. One hour after the final injection, mice were anesthetized with isoflurane and transcardially perfused with PBS containing 10 U/ml heparin, followed by 4% PFA. The dissected brains were fixed in 4% PFA for 24 h at 4 °C. Brains were then transferred to PBS containing 0.1% sodium azide until brain clearing and labeling. Brains were processed by LifeCanvas Technologies following the SHIELD protocol as previously published (*26, 71*). Samples were cleared for 7 days with Clear+ delipidation buffer, followed by batch labeling in SmartBatch+ with 8.6 μg Goat anti-GFP antibody (Encor GPCA-GFP), 17 μg anti-Mouse NeuN antibody (Encor MCA-1B7), and 6 μg anti-Rabbit FOS (Abcam ab214672) per brain. Fluorescently conjugated secondary antibodies were applied in 1:2 primary/secondary molar ratios (Jackson ImmunoResearch). Labeled samples were incubated in EasyIndex (LifeCanvas Technologies) for refractive index matching (n = 1.52) and imaged with SmartSPIM (LifeCanvas Technologies) at 4 μm z-step and 1.8 μm xy pixel size. Image analysis was conducted following the procedures as previously published (*26, 71*). Due to technical limitations raised by resolution limitations and distinct subcellular expression patterns of FOS (nuclear) and GFP (cytosolic) signals in three-dimensional space, the direct co-localization between FOS+ and GFP+ cells were not analyzed. For studies of acute exposure to and during spontaneous withdrawal from cocaine and morphine, brain were postfixed after perfusion as stated above, and serial 100um sections were collected between olfactory bulb and brainstem. FOS staining was conducted following standard immunohistochemistry procedures, and analysis was conducted using the open-source SMART pipeline as previously published (*72*).

### In silico FOS-Seq

FOS counts were z-scored within each batch of samples across conditions. Multiple batches of z-scored FOS activity were pooled for downstream analysis. After batch correction, images were co-registered with the Allen Brain Atlas reference mouse brain to demarcate region specificity of FOS signal. Next, normalization of FOS signal was achieved by subtracting average FOS levels in the saline condition from FOS levels in the cocaine and morphine conditions across brain regions. Region-specific gene expression data was then obtained from the Allen Brain Atlas *in situ* hybridization database, and brain areas identified to match between FOS imaging and in situ hybridization database were used for generating vectors for FOS and all identified genes. Each FOS vector and gene X ISH vector were first sigmoid transformed, followed by computations of Pearson Correlation Coefficients (PCCs) across brain regions for each condition. A histogram depicting the distribution of total PCCs was plotted. Next, a Gaussian distribution was fitted to this histogram (Fig. S5a, μ = 0.0, σ = 0.15). A threshold of ± 0.15, which corresponded to 1σ, was applied to identify genes that exhibited significant correlations. Genes correlated with PCC value > 0.15 or < −0.15 and p < 0.05 were classified as positively correlated or negatively correlated, and the rest were classified as not correlated. The method was adapted from a previously published approach (*26*). P-values are FDR-corrected at 5% threshold, referred to as adjusted P-values.

### snRNA-seq sample preparation

Animals were stereotaxically injected with AAV-Rheb-sgRNAs or AAV-scrambled-sgRNAs bilaterally in NAc core. To avoid confounds from Rheb and Control sgRNAs, each mouse was only transduced with one type of sgRNAs that was either Rheb-targeted or non-targeted without pooling AAVs. Three D1-Cas9 mice and three D2-Cas9 mice were transduced with Rheb-sgRNAs, and similarly three D1-Cas9 and three D2-Cas9 mice were transduced with Scrambled-sgRNAs. Three weeks after viral injections, mice were anesthetized under 5% isoflurane. NAc core was microdissected under a stereomicroscope in pre-chilled dissection buffer and immediately transferred to dry ice prior to downstream nuclei extraction. After dissection, frozen tissues on dry ice or stored in a −80°C freezer were immediately transferred to Teflon homogenizers containing 1 ml pre-chilled NP40 lysis buffer (Fisher Cat# FNN0021) and homogenized 15-30 times using a pellet pestle on ice.

Homogenized samples were incubated for another 10-15 min on ice, followed by passing through a 70 µm Flowmi Cell Strainer and a 40 µm Flowmi Cell Strainer (Millipore Sigma). The collected flowthrough was centrifuged at 500-1000 rcf for 5 min at 4°C and pellets were resuspended in staining buffer. 30% iodixanol buffer was then carefully loaded at the bottom of resuspended nuclei at 4°C. Samples were centrifuged at 10000 rcf for 20 min at 4°C. Supernatant containing debris was carefully removed. Pellets were resuspended in staining buffer containing anti-NeuN Alexa 647 antibody (abcam, Cat# ab190565) in order to enrich neurons. After antibody incubation and rotating for 30 min at 4°C, samples were washed with staining buffer without antibodies for 3 times. Samples were resuspended in FACS buffer after last-round wash and sent for FACS sorting. Hoechst 33342 (ThermoFisher Scientific, Cat# H3570) were added at a final concentration of 0.2 mM to label nuclei. Sorted nuclei were sent for downstream 10X genomics 5’ RNA-seq with CRISPR library preparation and sequenced using NovaSeq sequencer with ∼30000 reads/nuclei on average. Dissection buffer contains 1X HBSS, 2.5 mM HEPES-KOH [pH 7.4], 35 mM glucose, 4 mM NaHCO_3_, and actinomycin D (Sigma-Aldrich, Cat# A1410) at a final concentration of 20 µg/ml. NP40 lysis buffer contains 10 mM Tris-HCl [pH 7.4], 10 mM NaCl, 3 mM MgCl_2_, 0.1% NP40 dissolved in nuclease-free water. For 1 ml NP40 lysis buffer, 1 µl DTT, 25 µl 20 U/µl SupeRasine (Thermo Cat# AM2696), 12.5 µl 40 U/ µl RNasin (Promega Cat# N2615), 10 µl protease and phosphatase inhibitor cocktail (100X; Thermo Cat# 78442), 40 µl 1 mg/ml actinomycin D were added right before use. 30% iodixanol buffer contains 0.25 M sucrose, 25 mM KCl, 5 mM MgCl_2_, 20 mM Tricine-HCl [pH 8.0] and 30% Iodixanol dissolved in nuclease-free water. DTT, Superasine, Rnasin and protease inhibitors were added at the same concentration as NP40 lysis buffer right before use. Staining buffer contains 2% BSA, 0.05% NP40 dissolved in nuclease-free 1X PBS. Superasine, Rnasin and protease inhibitors were added at the same concentration as NP40 lysis buffer right before use. FACS buffer contains 2% BSA dissolved in nuclease-free 1X PBS buffer. Superasine, Rnasin and protease inhibitors were added at the same concentration as NP40 lysis buffer immediately prior to use.

### snRNA-seq analysis

Fastq files were aligned to mouse genome (mm10) and CRISPR sgRNAs feature sequences, and the expression levels in each cell were estimated with Cellranger (v 6.0.0). The gene expression count matrix for each sample was processed with the following steps: (1) Estimate doublet with Scrublet (https://github.com/swolock/scrublet) (*73*); (2) Estimate and correct the ambient RNA contaminations with SoupX (https://github.com/constantAmateur/SoupX) (*74*); (3) Load the corrected counting matrix into Seurat object with log normalization; (4) Calculate the proportion of UMIs from mitochondrial genes; (5) The cells assigned as doublets or mitochondrial content >1% were removed. The Seurat objects were integrated by following the RPCA workflow (https://satijalab.org/seurat/articles/integration_rpca.html) (*75*). The number of PCs used for UMAP calculation was selected with elbow plot (*76*). Then, the clustering was calculated with Leiden algorithm (*77*). Mixscape analysis was applied following the workflow (https://satijalab.org/seurat/articles/mixscape_vignette) (*38*). After Mixscape classification of perturbed vs. escaped cells, only perturbed cells and control cells were used for calculating differentially expressed genes. The differential gene expression of each comparison was performed with Seurat::FindMarkers() with logfc.threshold greater than 0.13, and raw p values from the returned set of genes were corrected using p.adjust() with the ‘BH’ method. Significant differential expression genes were defined as log2 (Fold Change) greater than 0.26 or less than −0.26, and corrected p values less than 0.05. GO analyses were conducted using the gseapy.enrichr() function in Python. Databases included GO_Biological_Process_2023, GO_Cellular_Component_2023, GO_Molecular_Function_2023. Enriched pathways with adjusted p values less than 0.05 were plotted.

### Chemogenetic modulations

5 mg/kg CNO was i.p. injected 20 min prior to injections of other solutions. For experiments aimed at activating retrograde NAc-projecting neurons, 2 mg/kg CNO was i.p. injected 30 min prior to providing access to food or water. Food consumption was measured in mice that had prior ad libitum access to food, 1 hour after the onset of the dark cycle, resembling a state of physiological hunger. Water consumption was measured 5 min after providing overnight water-deprived mice with free access to water.

### Optogenetic modulations

For photostimulating ChR2, a 473-nm laser (OEM Lasers/OptoEngine) was used to generate laser pulses (5-7 mW at the output tip of the fiber, 5 ms, 20 Hz) throughout the behavioral session, except when noted otherwise, controlled by a waveform generator (Keysight).

### Feeding and drinking behaviors

For optogenetic stimulation, mice were acclimated to a new clean cage for 5 minutes before experiments started. Mice were then provided free access to either food or water. For ChR2 stimulation in fasted or water-dehydrated mice, food or water intake was measured after 20 minutes of photostimulation, followed by 20 minutes with laser off.

### Open field assay

Mice were introduced into a 28 × 28 cm open field arena. Locomotor activity and fraction of movement over the 20-minute session were automatically tracked and quantified by Ethovision 9 (Noldus).

### Real-time place preference assay

Real-time place preference was conducted in a two-chamber acrylic box (50 × 25 × 25 cm). The preference between the two chambers was determined by the time spent on either side over a 20-minute session. The amount of time spent in each chamber was automatically tracked and calculated using the Ethovision 9 software (Noldus). Photostimulation of ChR2 (5-7 mW per fiber tip; 20 Hz; 10 ms pulses; 5 s on, 5 s off) was delivered by a mini-IO box (Noldus) when mice entered the designated light-paired side. Mice performed three test sessions with photostimulation.

### *In vivo* two-photon imaging

Two-photon (2P) calcium imaging was performed on a Scientifica SliceScope galvo-scanning 2P microscope with a Nikon 16×/0.8 water-dipping objective and a Coherent Chameleon Ultra II Ti:Sapphire laser source. The objective was focused onto the rear image plane of the implanted GRIN lens, so that the excitation laser beam was relayed into the sample by the GRIN lens, and conversely fluorescence was relayed out of the sample and recorded using photon-multiplier tubes in the SliceScope’s non-descanned detection head.

Microscope hardware and data acquisition was controlled using the ScanImage software (versions 5.5 and 5.6, Vidrio Technologies), which is based on Matlab (The Mathworks). The field-of-view of the microscope was sufficient to record the diameter of the GRIN lens image, 500 µm, at a frame rate of 4.82 Hz. Liquid food (Ensure) and water were provided via a spout connected to a touch detector (lickometer) during the imaging. Mice were adapted to liquid food as the only nutrition source for at least three days before the refeeding experiments. On experiment day, fasted or dehydrated mice went through a three-minute baseline recording, followed by 3 μL water or liquid food dispensed by the lickometer spout at the beginning of the consumption trial. After the trial started, the lickometer spout dispensed 3 μL water or liquid food upon each lick from mice.

### Drug administration assay

For behavioral experiments, a dosage of 20 mg/kg cocaine, or 10 mg/kg morphine dissolved in saline, respectively, was administered via daily single-dose i.p. injections. These doses were chosen based on their rewarding properties in conditioned place preference studies (*10*). Spontaneous withdrawal was defined as within 3 days after the last dose from chronic exposure of cocaine or morphine. During these 3 days, each cohort of mice received daily i.p. injections of saline. For *in vivo* two-photon calcium imaging experiments, a dosage of 10 mg/kg cocaine or 5 mg/kg morphine dissolved in saline was administered by single-dose i.p. injections. Notably, doses in the calcium imaging experiments were lowered to minimize excessive treadmill running which produce movement-related artefacts, while maintaining high enough doses that are known to be rewarding (*10, 78*). On the experimental day, mice were head-fixed and placed on the treadmill for 5 min habituation. Mice went through a one-minute baseline recording, followed by one i.p. injection of saline, cocaine or morphine. The neural responses of each mouse were then recorded for 1 minute every 10 minutes within one hour. All cohorts of mice received one daily i.p. injection of cocaine or morphine continuing for 5 days. Mice were able to freely walk on the treadmill, and velocity was recorded at the same time. Mice had *ad libitum* access to food and water before and after the imaging, but were restricted throughout experimental procedures.

### Data Analysis

Behavior data was analyzed using the Ethovision X9 software (Noldus). For two-photon imaging experiments, behavioral and imaging data was analyzed using the Suite2p pipeline (*79*) and custom Python scripts. Multiple trials within the same session were first concatenated, then processed by the Suite2p pipeline with non-rigid motion correction, to extract all neurons recorded across trials. According to the Suite2p pipeline, this concatenation does not normalize or alter the raw fluorescence of individual trials. Each session data from different days were processed independently. Neurons extracted by the pipeline were subsequently curated manually in the Suite2p graphical user interface. When aiming to compare responses of the same neurons across multiple sessions or days, spatial footprints of neurons from each session extracted by suite2p were mapped using CellReg (*80*). Consistent with prior reports (*81*), 20-50% of the same neurons were able to be tracked across multiple daily sessions.

To preprocess the data from food and water gel sensory responses, total fluorescence of each individual neuron was normalized using the formula z = (Fraw – μ)/σ, where Fraw is the raw fluorescence as extracted by the Suite2p pipeline, μ is the mean of Fraw during the baseline period (30 seconds prior to food and water gel presentation), and σ is the standard deviation of Fraw during the baseline period.

The data from liquid food and water consumption experiments was preprocessed as follows in order to reduce variabilities for multiple-day comparisons: The raw fluorescence was z-scored according to the formula z = (F– μ)/σ, where F is calculated from the raw fluorescence by applying a second order Butterworth filter with normalized cut-on frequency 0.266; μ is the mean baseline fluorescence (taken over the traces with activity below the median during the baseline period), σ is the standard deviation of fluorescence during the baseline period (taken over the traces with activity below the median during the baseline period). For this analysis, neurons with averaged responses larger than 3σ from 10 seconds after consumption start were considered to be activated.

Clustered neural traces during consumption were identified by k-means clustering. The neuronal states based on the k-means clustering label were projected to two-dimensional space using the t-SNE algorithm or non-negative matrix factorization (NMF) for visualization.

To preprocess the data from the drugs of abuse experiments, the raw fluorescence traces were filtered as described above. Within each session, each 1-minute imaging trace was normalized as Fcorr = (F– μ)/ μ, where F is the raw fluorescence and μ is the average of the lowest quintile of the raw fluorescence. All corrected traces were concatenated and used for downstream analyses. Peaks of neuronal responses were identified using the SciPy package (find_peak() function) with the peak height threshold set to 3σ) and minimum distance between peaks set to ∼1.3 seconds (based on the GCaMP6s decay time). Strength of neuronal preference was defined as: (Peak_cocaine_ – Peak_food/water_)/(Peak_cocaine_ + Peak_food/water_). To quantify cocaine-activated neurons (peak greater than 3σ), the timespan from 20-40 minutes post i.p. injection was used, while 30-50 minutes post i.p. injection was used to quantify morphine-activated neurons. The timespan was chosen based on observed behavioral effects (Fig. S1), neuronal dynamics (Fig. S4), and established pharmacokinetics from literature for cocaine/morphine post i.p. injections (*82, 83*).

To differentiate the motor-associated from the non-motor-associated neural correlates, we first converted the animals’ walking velocity into binary movement vectors (moving vs. not moving) as recorded from the treadmill and then computed the cross-correlation (evaluated for a range of 1-10 frames lag, corresponding to ∼2 seconds) between neuronal traces and binary movement vectors (as computed by thresholding the raw velocity measurement vector). The lag with the highest cross-correlation was chosen for downstream analyses. We next computed the Pearson correlation coefficients (PCC) between the lag-corrected neuronal traces and the binary movement vector. We selected highly motion-correlated neuronal traces as those with the modulus of their PCC with the movement vector above a threshold in the range 0.2-0.4. We adjusted the threshold based on the principle that the averaged frames of neural activities (above median) during movement periods are no less than those during the non-movement period, Wilcoxon test was conducted to compare the significant difference level between the movement versus non-movement frames. The neurons showing PCCs larger than the threshold were considered to be motor-associated neurons, the rest were nonmotor-associated neurons. Each neuronal pair with activity correlation coefficient greater than 0.3 was considered to be “synchronized”. A connectivity index was then calculated as the total number of pairs of synchronized neurons divided by the pairs of synchronized neurons during the baseline period (before i.p. injection).

Tensor component analysis (TCA) was conducted as described previously (*25*). The tensor matrix comprised of trial, neuron and time series of neuronal activity was loaded into the TCA algorithm. The factor matrices were constrained to be nonnegative. Each neural state was averaged from multiple sessions recorded from each cohort of mice. Neurons positively contributing to state 1 relative to state 2 were determined by positive values after the subtraction: state 1 loadings – state 2 loadings.

### Statistics and reproducibility

Statistical analyses were conducted using Graphpad Prism 9.5. Throughout the paper, values are reported as mean ± s.e.m. (error bar or shaded area). P-values for pair-wise comparisons were obtained using the two-tailed Wilcoxon signed-rank test. P-values for comparisons across multiple groups were conducted using ANOVA (with repeated measures when possible) and corrected for multiple comparisons. The statistical models used for imaging data analysis as described above were carried out using the scikit-learn Python package (*84*). No statistical methods were used to predetermine sample size. The experiments were not randomized. The investigators were not blinded to allocation during experiments and outcome assessment. Representative images were selected from 3 to 5 original biological replicates.

---

**Figure. S1.**
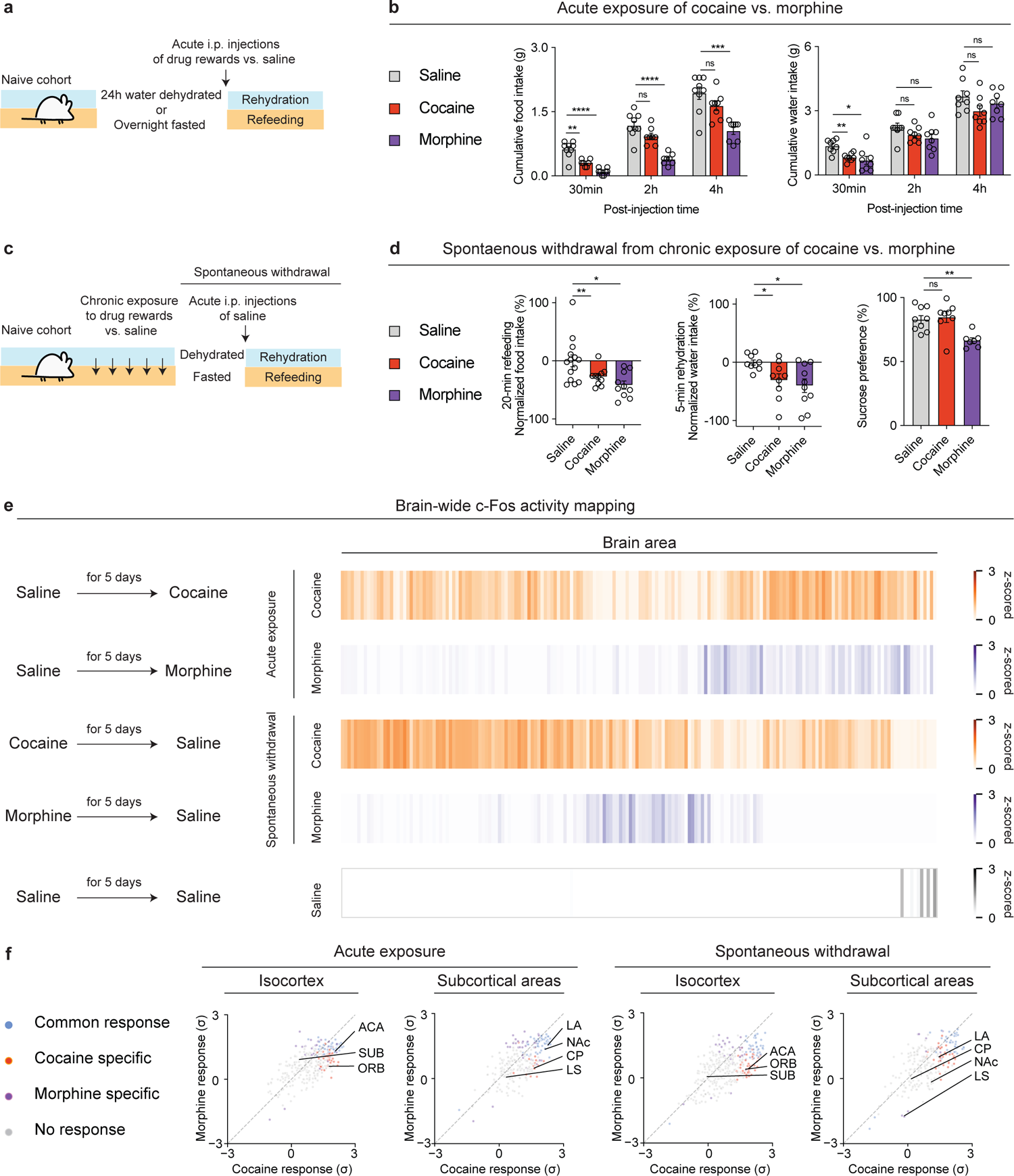
Acute exposure and spontaneous withdrawal from cocaine vs. morphine both decrease consumption of natural rewards while activating common vs. distinct brain areas. (a), Schematic of the experimental design for refeeding and water rehydration assays post acute exposure of 20 mg/kg cocaine, 10 mg/kg morphine vs. saline. Comparisons of (b), Cumulative food intake in fasted mice (left panel), Cumulative water intake in water-dehydrated mice (right panel) received i.p. injections of saline, cocaine, morphine 20 min prior to providing free access to food or water (n = 9, 8, 8 for each group, respectively, two-way ANOVA with Dunnett’s multiple comparisons). (c), Schematic of the experimental design of refeeding and water rehydration assays during spontaneous withdrawal from chronic exposure of 20 mg/kg cocaine, 10 mg/kg morphine vs. saline. Comparisons of (d), Normalized food intake in refeeding assay from fasted mice during spontaneous withdrawal from saline, cocaine, morphine (n = 14, 10, 10 for each group, one-way ANOVA with Dunnett’s T3 multiple comparisons, data were pooled from 2 cohorts, left panel). Normalized water intake in rehydration assay from water-dehydrated mice during spontaneous withdrawal from saline, cocaine, morphine (n = 9, 9, 10 for each group, data were pooled from 3 cohorts, one-way ANOVA with Dunnett’s T3 multiple comparisons, data were pooled from 2 cohorts, middle panel). Comparisons of sucrose preference vs. water (%) in water-dehydrated mice during spontaneous withdrawal (n = 9, 8, 7 for saline, cocaine, morphine group, respectively, one-way ANOVA with Dunnett’s T3 multiple comparisons, right panel). (e), Heatmap overview of brain areas showing significantly differential FOS activity in response to acute 20 mg/kg cocaine, acute 10 mg/kg morphine, spontaneous withdrawal from chronic exposure of cocaine and morphine vs. saline (n = 3, 4, 4, 4, 5 for each group, one-way ANOVA for each brain area with cut-off p < 0.05 classified as statistically significant, followed by K-means clustering). (f), Scatter plot of FOS activity after acute exposure to cocaine vs. morphine (left). Scatter plot of FOS activity of after spontaneous withdrawal from cocaine vs. morphine (right). All error bars represent mean ± s.e.m. NS, not significant, *P < 0.05, **P < 0.01, ***P < 0.001, ****P < 0.0001.

**Figure. S2.**
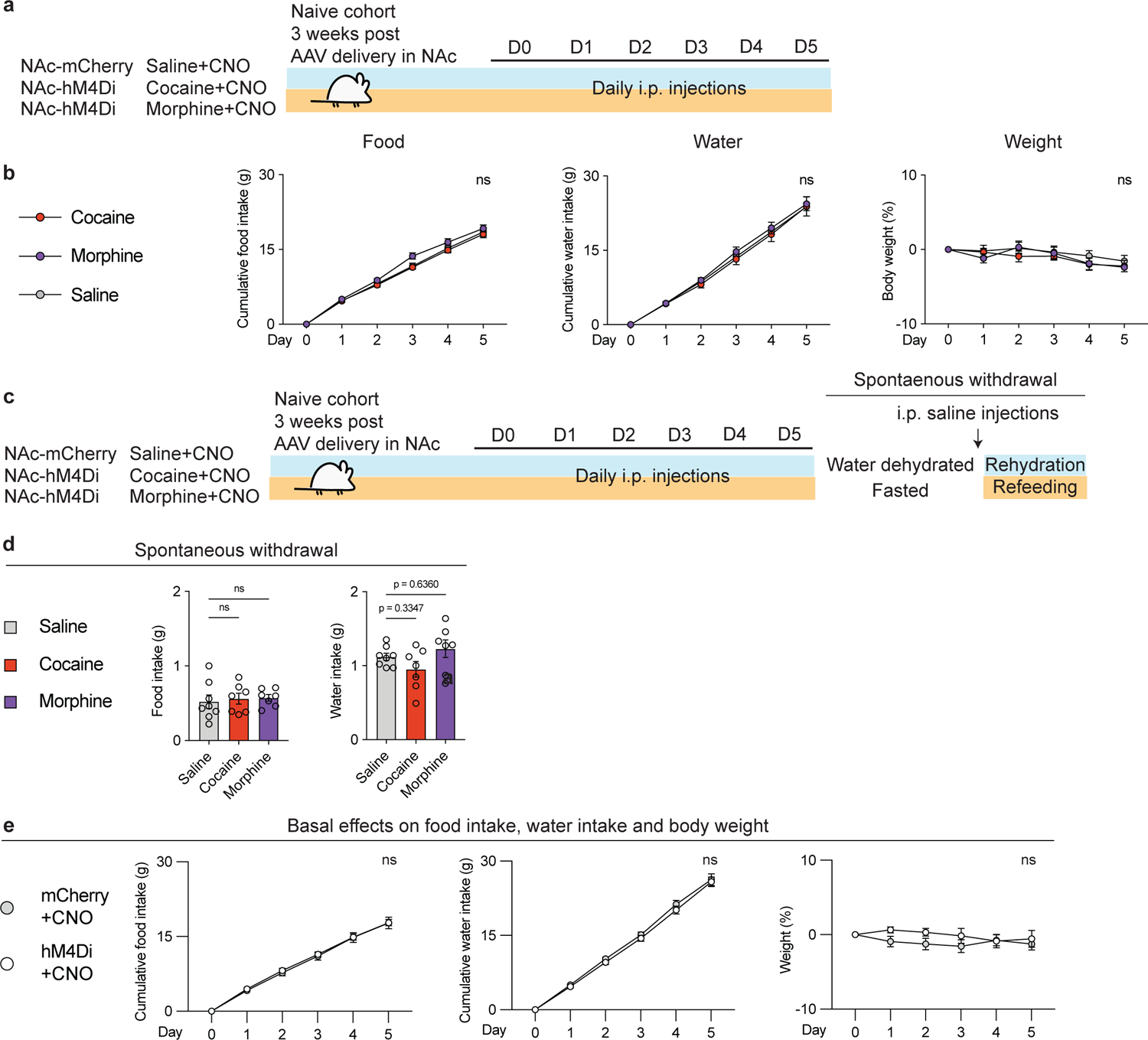
Chemogenetic silencing of NAc neurons prevents attenuated consumption of food and water specifically induced by chronic exposure to cocaine or morphine. (a), Schematic of the experimental design for chemogenetic modulations of NAc neurons. Three cohorts of mice respectively expressing mCherry, hM4Di or hM4Di in NAc received 5 mg/kg CNO i.p injections 20 min prior to receiving drug rewards vs. saline for 5 days. (b), Comparisons of Cumulative food intake (g), Cumulative water intake (g), Weight (%) in NAc-mCherry + saline + CNO, NAc-hM4Di + 20 mg/kg cocaine + CNO, NAc-hM4Di + 10 mg/kg morphine + CNO, respectively (n = 8, 7, 7 for each group, two-way ANOVA with Dunnett’s multiple comparisons). (c), Schematic of the experimental design for the chemogenetic silencing. Three cohorts of mice expressing mCherry, hM4Di or hM4Di, respectively, received daily i.p. injections of 5 mg/kg CNO 20 min prior to receiving i.p. injections of saline, 20 mg/kg cocaine, 10 mg/kg morphine for 5 days. (d), Comparisons of Food intake (left panel), Water intake (right panel) in fasted mice or water-dehydrated mice during spontaneous withdrawal from saline, cocaine, morphine (n = 8, 7, 7 for each group, one-way ANOVA with Dunnett’s T3 multiple comparisons). (e), Comparisons of Cumulative food intake, Cumulative water intake, Weight of mice expressing mCherry vs. hM4Di received daily i.p. injections of 5 mg/kg CNO for 5 days (n = 8, 7 for each group, two-way ANOVA with Šídák’s multiple comparisons). All error bars represent mean ± s.e.m. NS, not significant, *P < 0.05, **P < 0.01, ***P < 0.001, ****P < 0.0001.

**Figure. S3.**
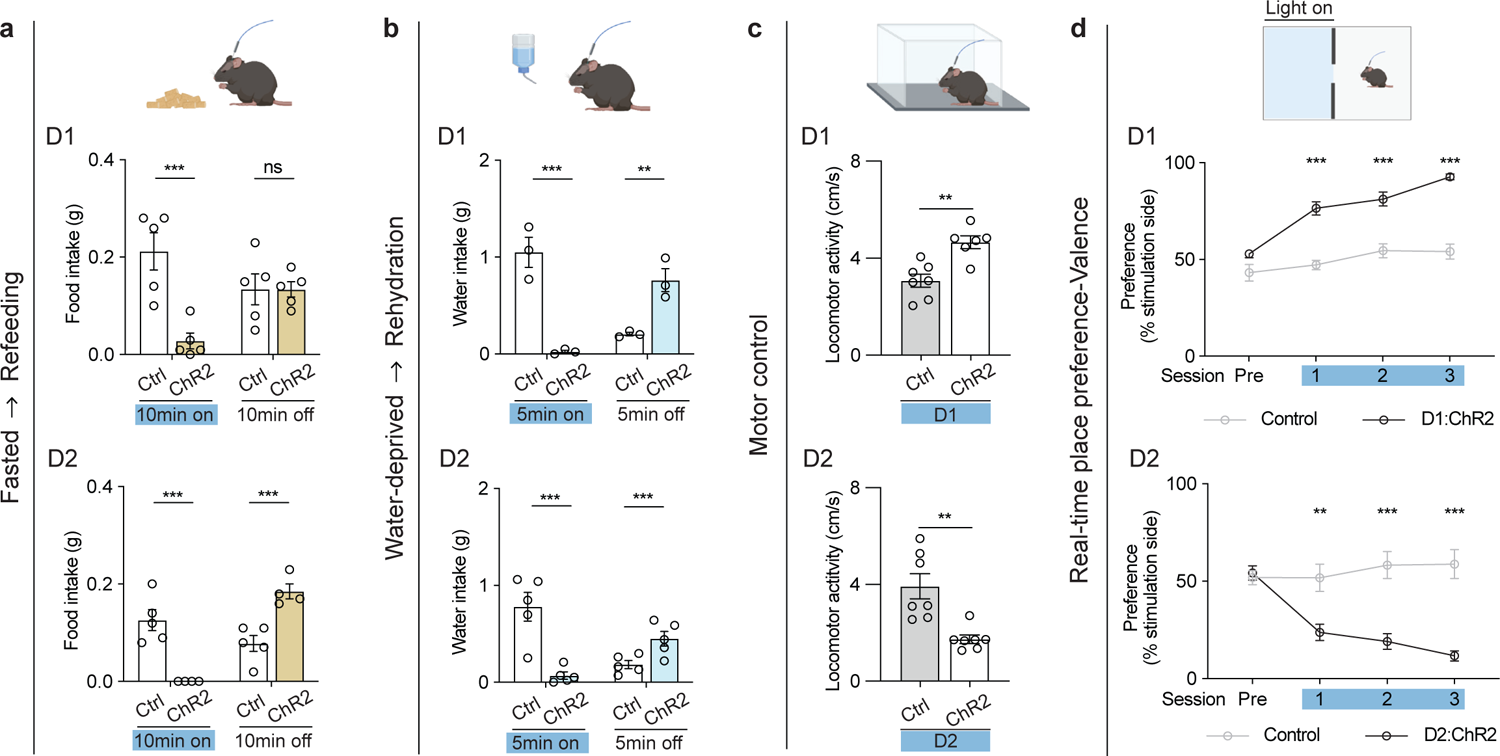
Optogenetic activation of NAc neurons regulates key aspects of natural reward processing. (a), Schematic of the refeeding assay in fasted mice that underwent optogenetic activations of D1^NAc^ or D2^NAc^ neurons vs. the control groups. Comparisons of food intake in Control vs. D1-ChR2 group (n = 5, 5 for each group, two-way ANOVA with Šídák’s multiple comparisons, top panel). Comparisons of food intake in Control vs. D2-ChR2 group (n = 5, 4 for each group, two-way ANOVA with Šídák’s multiple comparisons, bottom panel). (b), Schematic of the rehydration assay in water-dehydrated mice that underwent optogenetic activations of D1^NAc^ or D2^NAc^ neurons vs. the control groups. Comparisons of water intake in Control vs. D1-ChR2 group (n = 3, 3 for each group, two-way ANOVA with Šídák’s multiple comparisons, top panel). Comparisons of water intake in Control vs. D2-ChR2 group (n = 5, 5 for each group, two-way ANOVA with Šídák’s multiple comparisons, bottom panel). (c), Schematic of the open field test in ad libitum fed mice that underwent optogenetic activations of D1^NAc^ or D2^NAc^ neurons vs. the control groups. Comparisons of locomotor activity in the Control vs. D1-ChR2 group (n = 7, 6 for each group, two-tailed Student’s t tests, top panel). Comparisons of locomotor activity in the Control vs. D2-ChR2 group (n = 7, 7 for each group, two-tailed Student’s t tests, bottom panel). (d), Schematic of real-time place preference assay in ad libitum fed mice that underwent optogenetic activations of D1^NAc^ or D2^NAc^ neurons vs. the control groups. Comparisons of the preference at the stimulation side in the Control vs. D1-ChR2 group (n = 5, 5 for each group, two-way ANOVA with Šídák’s multiple comparisons, top panel). Comparisons of the preference at the stimulation side in the Control vs. D2-ChR2 group (n = 5, 6 for each group, two-way ANOVA with Šídák’s multiple comparisons, bottom panel). Conditions where mice received 473 nm laser on are highlighted in blue. All error bars represent mean ± s.e.m. NS, not significant, *P < 0.05, **P < 0.01, ***P < 0.001, ****P < 0.0001.

**Figure. S4.**
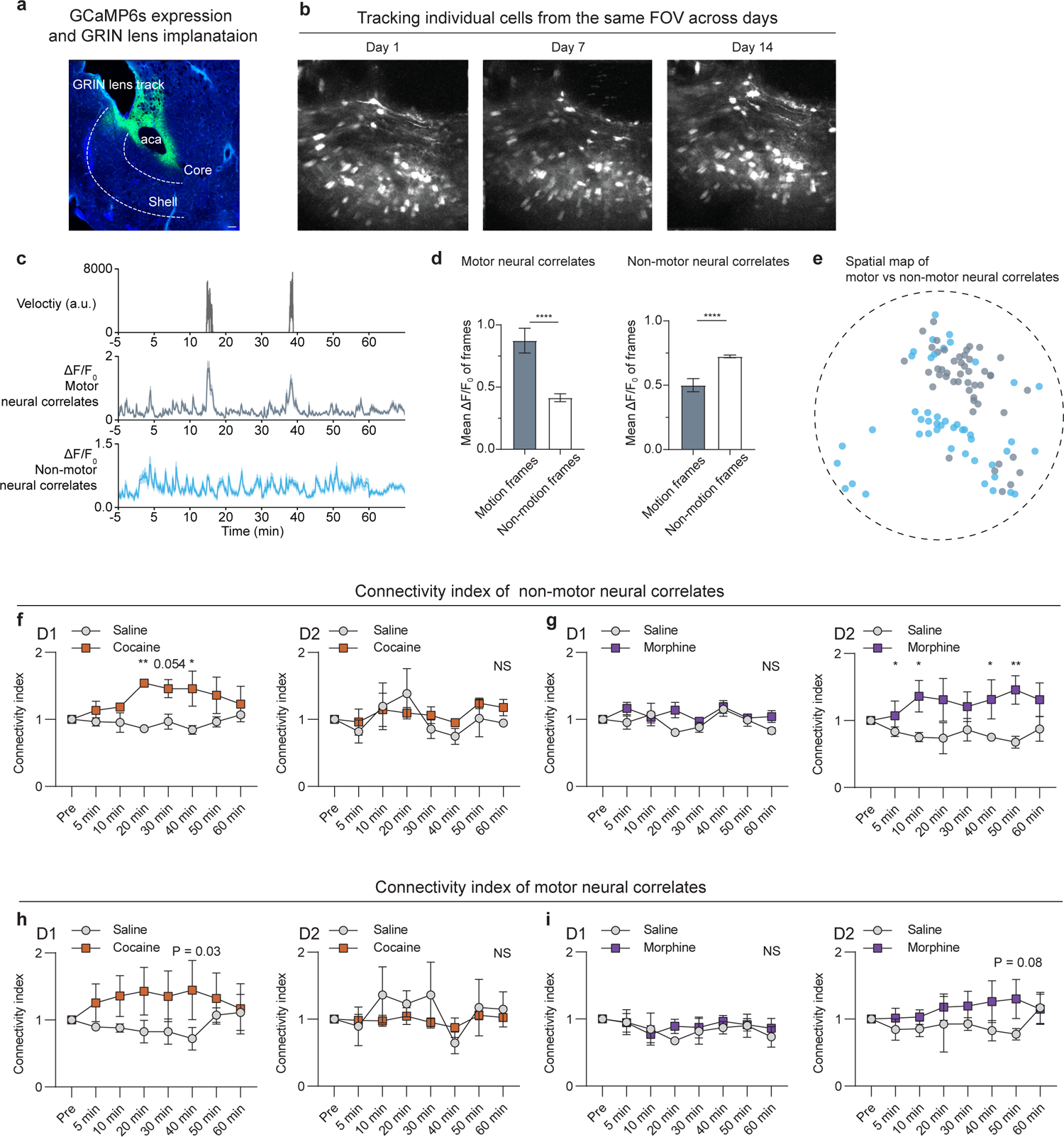
Cocaine vs. morphine selectively synchronizes NAcc^D1^ or NAcc^D2^ neuronal firing pattern, especially on nonmotor-associated neurons. (a), Example image of GCaMP6s expression in the NAc core after viral transduction. (b), Example field of views (FOV) across days. (c), Example velocity traces vs. averaged neural traces of motor-vs. nonmotor-associated neurons. (d), Example comparison of movement frames vs. nonmovement frames of motor-(n = 39 neurons) vs. nonmotor-associated (n = 92 neurons) neurons (two-tailed Wilcoxon test). (e), Example spatial distribution of motor-vs. nonmotor-associated neurons in the imaging focal plane. (d), Quantification of connectivity index averaged from all sessions of D1 nonmotor neural correlates modulated by cocaine compared to saline (n = 3 mice, two-way ANOVA, with Sidak’s multiple comparisons). (left panel), Quantification of connectivity index averaged from all sessions of D2 nonmotor neural correlates modulated by cocaine compared to saline (n = 3 mice, two-way ANOVA, with Sidak’s multiple comparisons) (right panel). (f), Quantification of connectivity index averaged from all sessions of D1 nonmotor neural correlates modulated by morphine compared to saline (n = 3 mice, two-way ANOVA, with Sidak’s multiple comparisons, left panel), Quantification of connectivity index averaged from all sessions of D2 nonmotor neural correlates modulated by morphine compared to saline (n = 3 mice, two-way ANOVA, with Sidak’s multiple comparisons, right panel). (g), Quantification of connectivity index averaged from all sessions of D1 motor neural correlates modulated by cocaine compared to saline (n = 3 mice, two-way ANOVA, with Sidak’s multiple comparisons, left panel). Quantification of connectivity index averaged from all sessions of D2 motor neural correlates modulated by cocaine compared to saline (n = 3 mice, two-way ANOVA, with Sidak’s multiple comparisons, right panel). (h), Quantification of connectivity index averaged from all sessions of D1 motor neural correlates modulated by morphine compared to saline (n = 3 mice, two-way ANOVA, with Sidak’s multiple comparisons, left panel). (i), Quantification of connectivity index averaged from all sessions of D2 motor neural correlates modulated by morphine compared to saline (n = 3 mice, two-way ANOVA, with Sidak’s multiple comparisons, right panel). All error bars represent mean ± s.e.m. NS, not significant, *P < 0.05, **P < 0.01, ***P < 0.001, ****P < 0.0001.

**Figure. S5.**
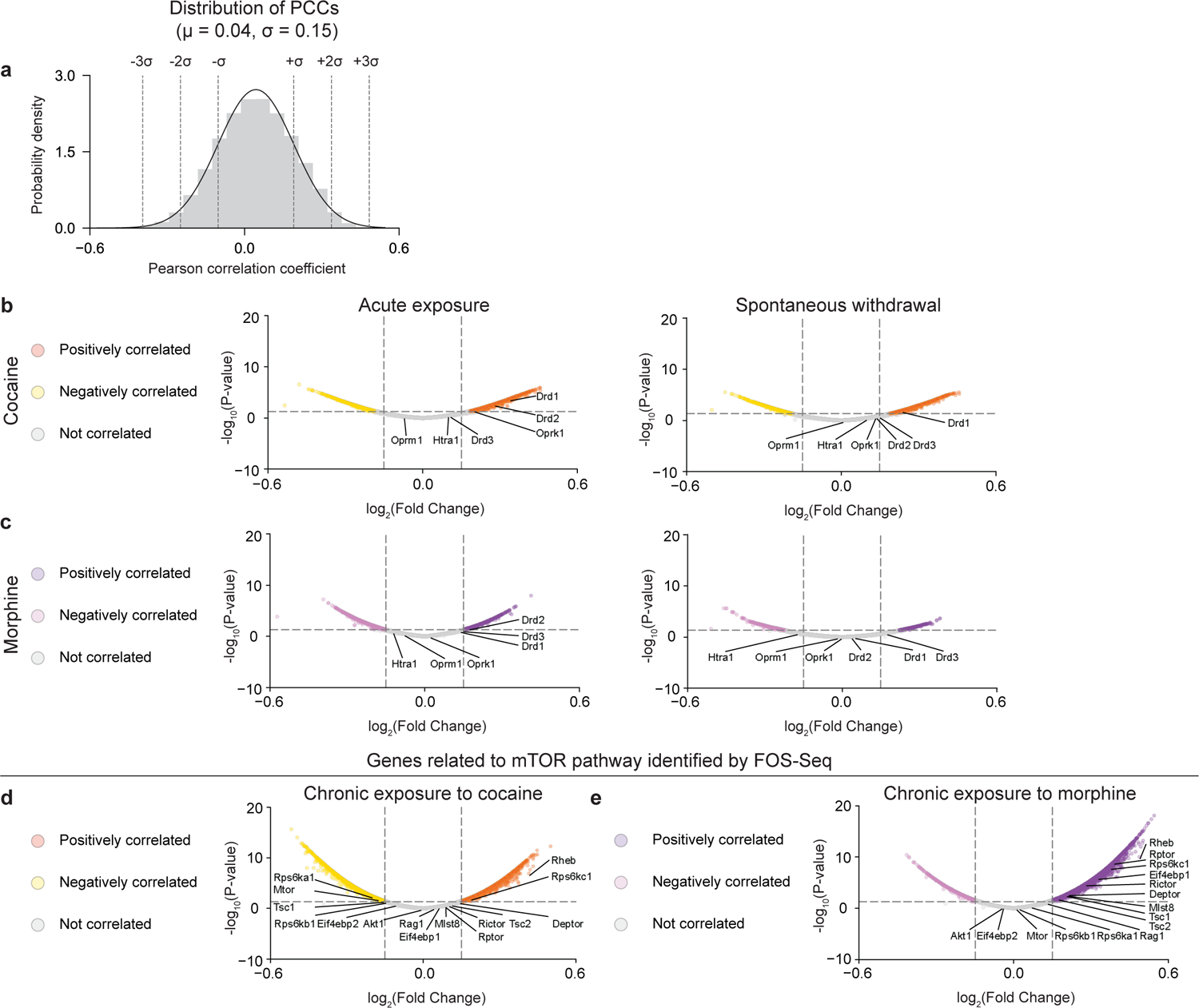
*In silico* FOS-Seq approach identifies canonical marker genes associated with acute exposure and spontaneous withdrawal from cocaine vs. morphine. (a), Histogram distribution of Pearson correlation coefficients (PCC) pooled from all the data after FOS-Seq. Volcano plot of genes correlated with (b), acute exposure of cocaine (left panel), spontaneous withdrawal from cocaine (right panel). (c), acute exposure of morphine (left panel), spontaneous withdrawal from morphine (right panel). (d), Volcano plot of genes in the mTOR pathway correlated with chronic exposure of cocaine. (e), Volcano plot of genes in the mTOR pathway correlated with chronic exposure of morphine. Genes with Pearson Correlation Coefficient > 0.15 or < −0.15, and p < 0.05 are classified as Positively correlated or Negatively correlated, the rest are classified as Not correlated.

**Figure. S6.**
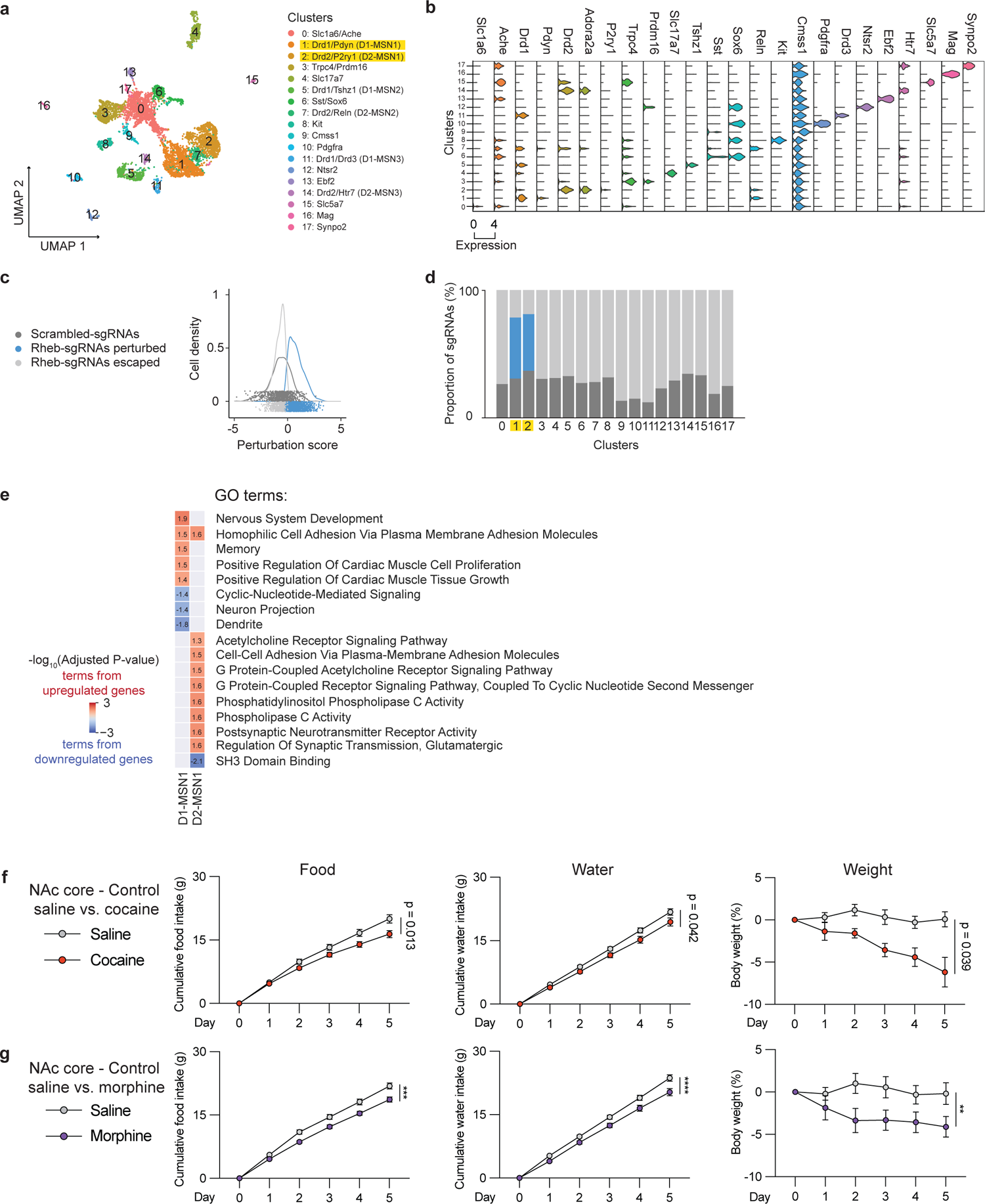
snRNA-seq after CRISPR perturbation of *Rheb* reveals cell-type-specific transcriptional regulation. (a), UMAP distribution of 18 clusters. (b), Stacked violin plot of marker gene expressions for each cluster. (c), Distribution of individual cell’s perturbation scores. (d), Composition of Rheb-sgRNAs perturbed, Rheb-sgRNAs escaped and the control Scrambled-sgRNAs across each cluster. (e), Significant GO terms from the differentially expressed genes of D1-MSN1 vs. D2-MSN1 (Adjusted P-value < 0.05). (f), Comparisons of cumulative food intake (g), water intake (g), weight (%) in the Control group treated with saline for 5 days followed by another 5-day cocaine treatment (n = 8, 8 for each group, two-way ANOVA w’th Šídák’s multiple comparisons). (g), Comparisons of cumulative food intake (g), water intake (g), weight (%) in the Control group treated with saline for 5 days followed by another 5-day morphine treatment (n = 8, 8 for each group, two-way ANOVA w’th Šídák’s multiple comparisons). All error bars represent mean ± s.e.m. NS, not significant, *P < 0.05, **P < 0.01, ***P < 0.001, ****P < 0.0001.

**Figure. S7.**
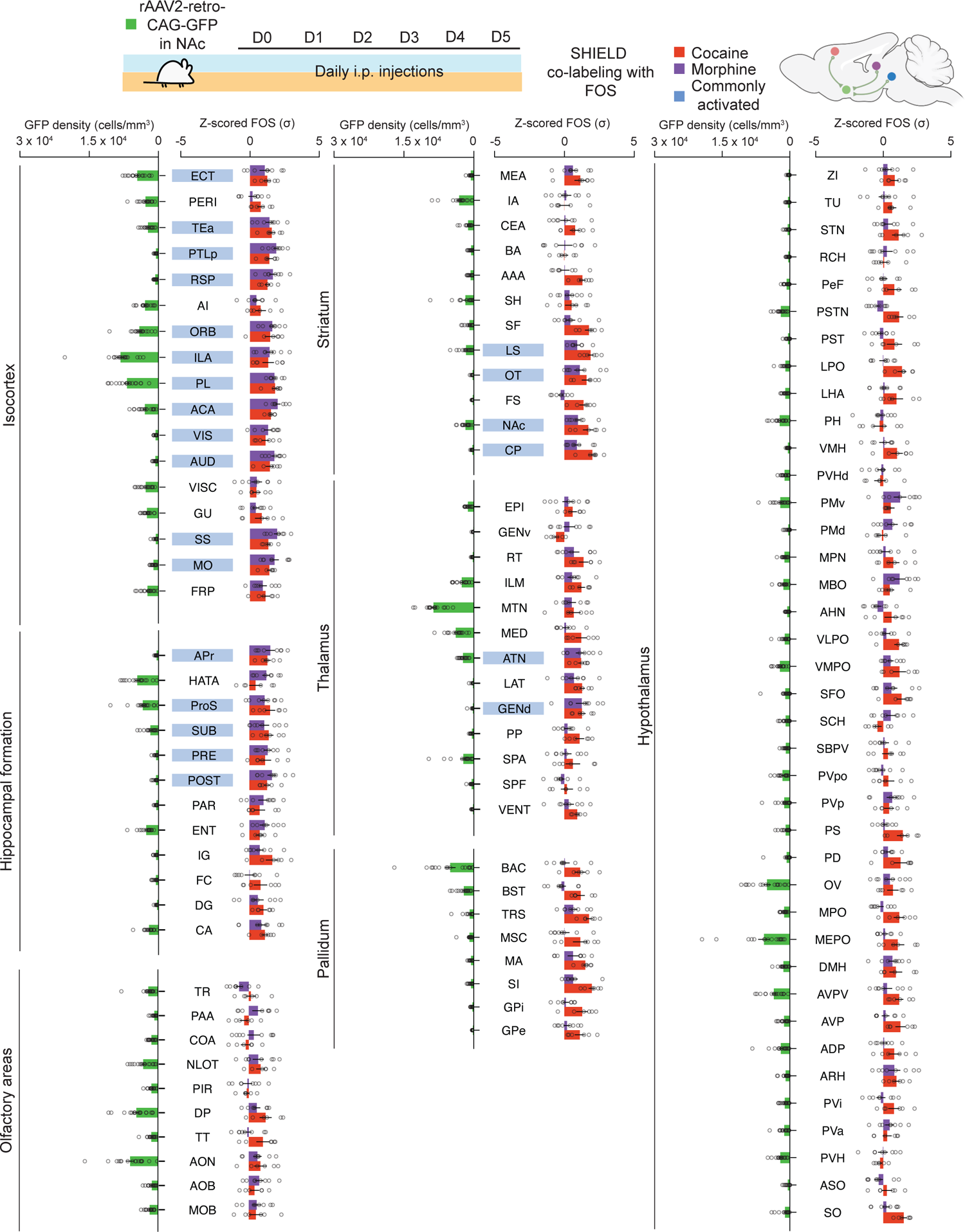
Simultaneous FOS activity mapping and monosynaptic retrograde tracing from NAc in response to cocaine vs. morphine. Schematic of experimental design for the retrograde tracing from NAc and chronic exposure of drug rewards. Three cohorts of mice with rAAV2-retro-CAG-GFP delivered in NAc received daily i.p. injections of 20 mg/kg cocaine, 10 mg/kg morphine and saline for 5 days (Top panel). GFP densitiy and FOS activity in each brain area (n = 17 for GFP density pooled from n = 6 mice received saline, n = 6 mice received cocaine, n = 5 mice received morphine, FOS activity is pooled from multiple batches and normalized by z-score with n = 6 mice received cocaine, n = 8 mice received morphine followed by subtraction of averaged z-score from n = 9 saline control, bottom panel). All error bars represent mean ± s.e.m. NS, not significant, *P < 0.05, **P < 0.01, ***P < 0.001, ****P < 0.0001.

**Figure. S8.**
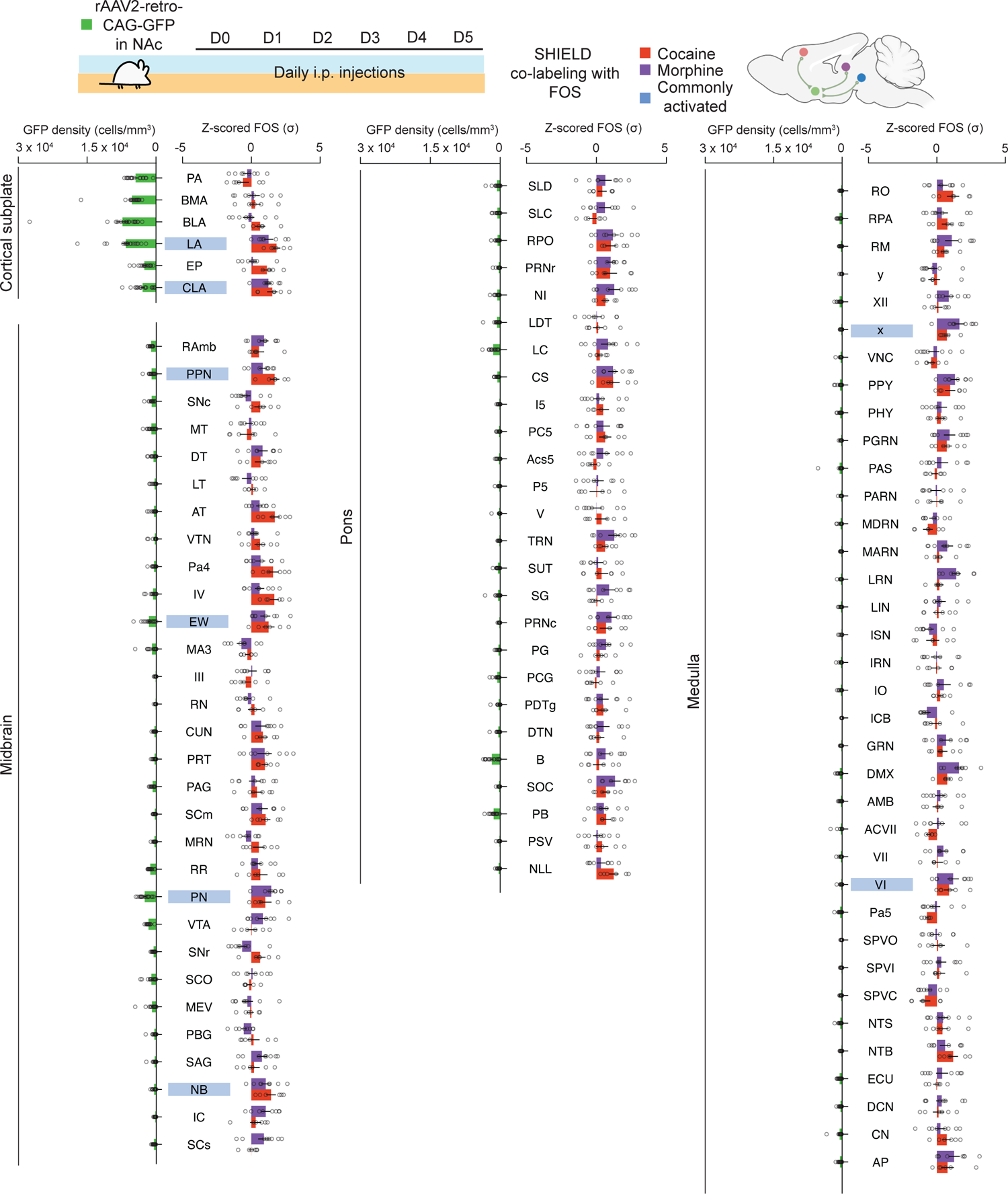
(Continued from Figure S7) Simultaneous FOS activity mapping and monosynaptic retrograde tracing from NAc in response to cocaine vs. morphine. Schematic of experimental design for the retrograde tracing from NAc and chronic exposure of drug rewards. Three cohorts of mice with rAAV2-retro-CAG-GFP delivered in NAc received daily i.p. injections of 20 mg/kg cocaine, 10 mg/kg morphine and saline for 5 days (top panel). GFP densitiy and FOS activity in each brain area (n = 17 for GFP density pooled from n = 6 mice received saline, n = 6 mice received cocaine, n = 5 mice received morphine, FOS activity is pooled from multiple batches and normalized by z-score with n = 6 mice received cocaine, n = 8 mice received morphine followed by subtraction of averaged z-score from n = 9 saline control, bottom panel). All error bars represent mean ± s.e.m. NS, not significant, *P < 0.05, **P < 0.01, ***P < 0.001, ****P < 0.0001.

**Figure. S9.**
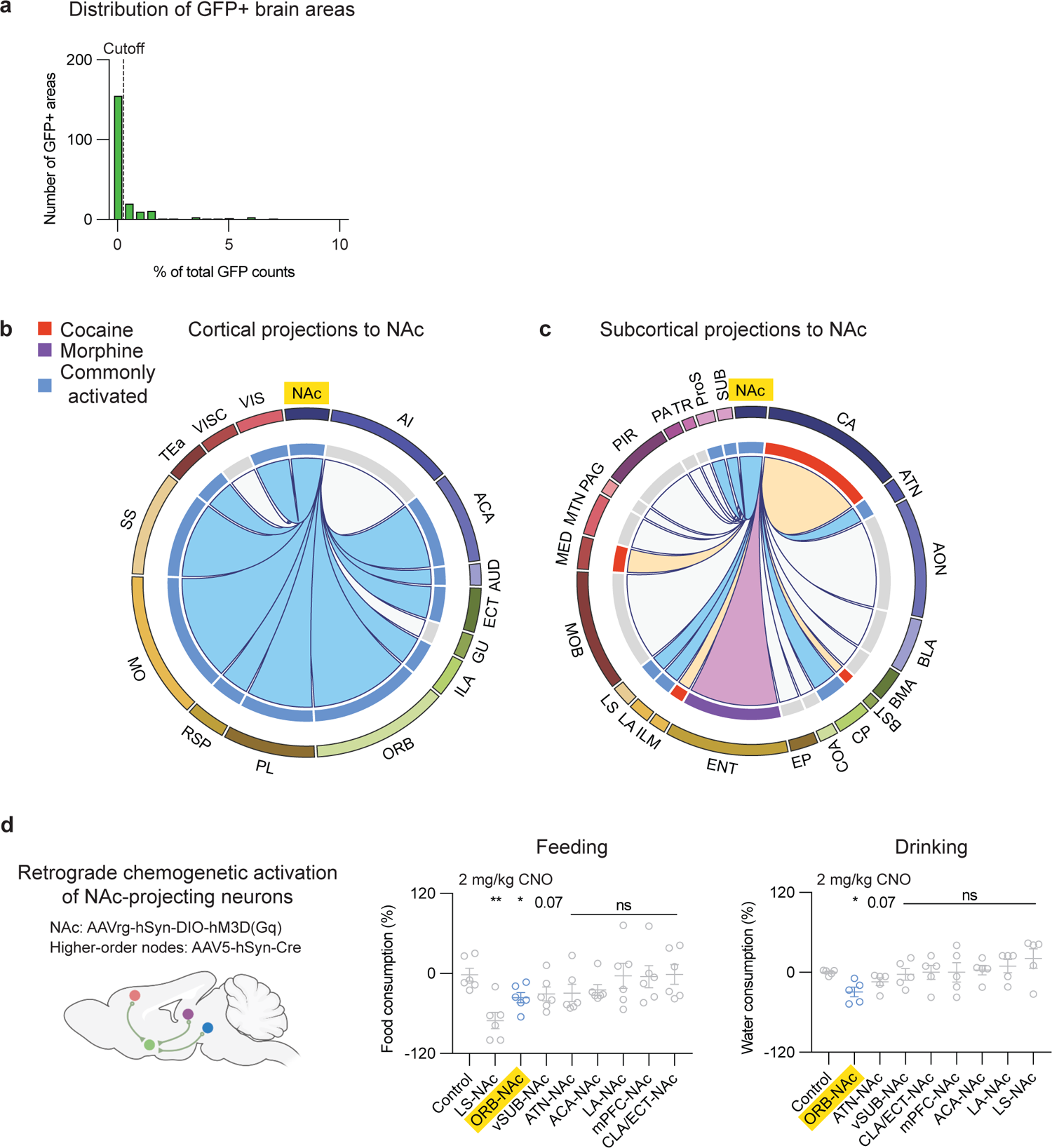
Brain-wide distribution of cocaine- and morphine-activated brain areas projecting to NAc. (a), Histogram distribution of the number GFP+ brain areas vs. percentage of total GFP counts. GFP+ brain areas with %GFP counts > 0.5% are used for generating the following circos maps (b), A circos map summarizing cortical projections to the NAc. (c), A circos map summarizing subcortical projections to the NAc. (d), Retrograde chemogenetic activation of NAc-projecting neurons in each higher-order node identified from the above circos maps. Food or water consumption was measured 30 min after i.p. injections of 2 mg/kg CNO. (n = 5 for each group of mice, Welch’s ANOVA test). All error bars represent mean ± s.e.m. NS, not significant, *P < 0.05, **P < 0.01, ***P < 0.001, ****P < 0.0001.

## Notes

### Competing Interest Statement

The authors have declared no competing interest.

